# In-cell architecture of an actively transcribing-translating expressome

**DOI:** 10.1101/2020.02.28.970111

**Authors:** Francis J. O’Reilly, Liang Xue, Andrea Graziadei, Ludwig Sinn, Swantje Lenz, Dimitry Tegunov, Cedric Blötz, Wim J. H. Hagen, Patrick Cramer, Jörg Stülke, Julia Mahamid, Juri Rappsilber

**Author notes:** These authors contributed equally to this work (Francis J. O’Reilly, Liang Xue, Andrea Graziadei).

## Abstract

Structural biology performed inside cells can capture molecular machines in action within their native context. Here we develop an integrative in-cell structural approach using the genome-reduced human pathogen *Mycoplasma pneumoniae*. We combine whole-cell crosslinking mass spectrometry, cellular cryo-electron tomography, and integrative modeling to determine an in-cell architecture of a transcribing and translating expressome at sub-nanometer resolution. The expressome comprises RNA polymerase (RNAP), the ribosome, and the transcription elongation factors NusG and NusA. We pinpoint NusA at the interface between a NusG-bound elongating RNAP and the ribosome, and propose it could mediate transcription-translation coupling. Translation inhibition dissociates the expressome, whereas transcription inhibition stalls and rearranges it, demonstrating that the active expressome architecture requires both translation and transcription elongation within the cell.

## Main Text

The two fundamental processes of gene expression, transcription and translation, are functionally coupled in bacterial cells. While the transcribing RNA polymerase (RNAP) produces a nascent mRNA chain that can be directly translated by ribosomes (*1*–*3*), translation has been shown to influence the overall transcription rate in *Escherichia coli* (*4, 5*). Recently, the structure of an *in vitro*-reconstituted RNAP-ribosome supercomplex was reported and referred to as “expressome” (*6*). Another structure of RNAP in complex with the small ribosomal subunit showed a different arrangement (*7*). These structures imply a physical link between the two processes. However, neither reconstitution included NusG, which was previously proposed to link RNAP and the ribosome (*8, 9*), or other essential transcription elongation factors (*10*), raising questions with regard to the context-distinct mechanisms of coupling that could be utilized *in vivo*.

To structurally analyse transcription-translation coupling inside cells, we combined in-cell crosslinking mass spectrometry (CLMS) (*11*) and cellular cryo-electron tomography (cryo-ET) (*12*). We used the small, genome-reduced bacterium *Mycoplasma pneumoniae*, which is an ideal cell model for system-wide structural studies (*13*). While the *M. pneumoniae* genome has undergone significant reduction during evolution as a human pathogen, it has retained the core transcription and translation machineries (*14*–*16*). Furthermore, RNAP and the ribosome in *M. pneumoniae* have been suggested to form a supercomplex by tandem affinity-purification mass spectrometry (*13*).

To assess the topology of this putative RNAP-ribosome supercomplex and their associated regulatory factors, we performed whole-cell CLMS by adding the membrane-permeable crosslinkers disuccinimidyl suberate (DSS) and disuccinimidyl sulfoxide (DSSO) to *M. pneumoniae* cells (*11*) (fig. S1 and table S1). These irreversibly crosslink nucleophilic side chains of amino acid residues found within 11.4 Å and 10.1 Å, respectively. Using xiSEARCH (*17*) and xiFDR (*18*), we identified 10,552 intra-protein crosslinks (unique residue pairs) and 1,957 inter-protein crosslinks with a 5% crosslink-level false discovery rate (FDR), representing 577 distinct protein-protein-interactions (PPIs) at a 5% PPI-level FDR (Fig. 1A and supplementary text). Identified crosslinks cover 83% of the detectable proteome (table S2 and fig. S1) and identify PPIs involving membrane proteins (41 % of all detected PPIs), as well as 76 currently uncharacterised proteins (fig. S1-S2). The ribosome, RNAP and associated factors make up some of the most abundant complexes in *M. pneumoniae* cells, and were consequently identified with several unique crosslinks, providing information on the structural organization of their interactions (Fig. 1B and fig. S2- S3).

**Fig. 1.**
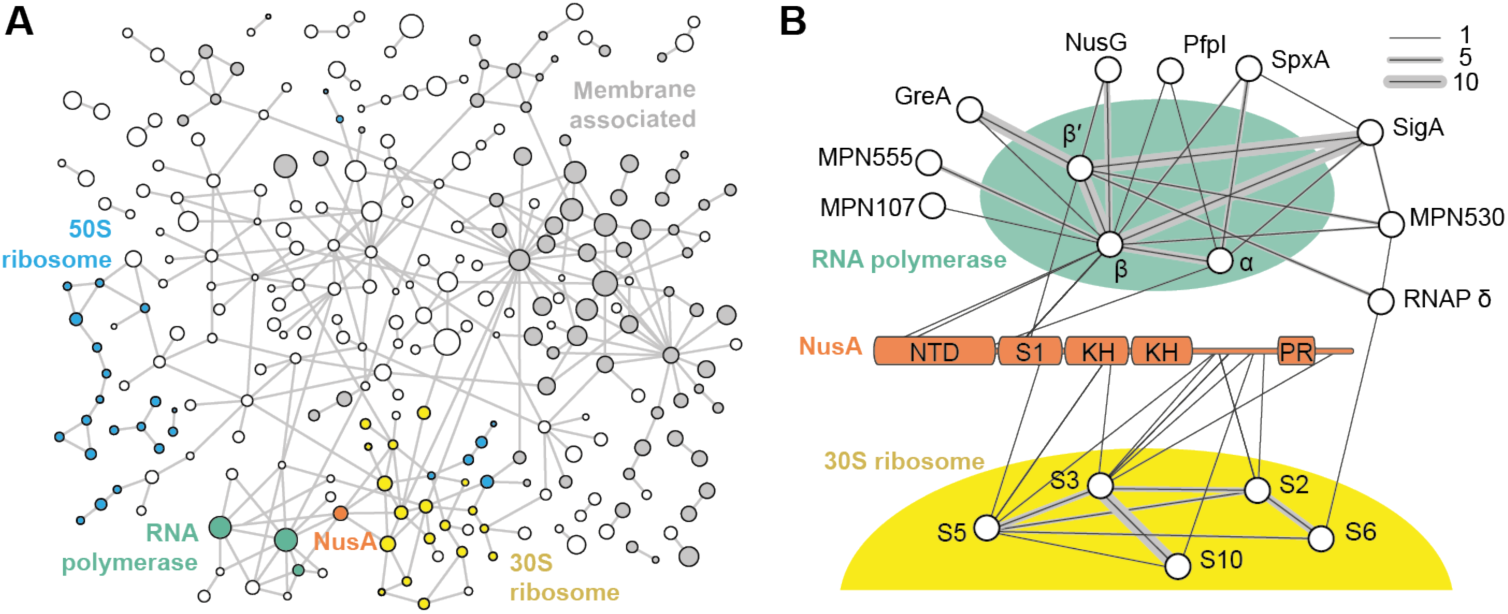
Crosslink-based protein interaction map of *M. pneumoniae* proteome. (**A**) 577 distinct protein-protein interactions at 5% PPI-level FDR are identified (interactions to 8 chaperones and highly abundant glycolytic enzymes are removed for clarity). Membrane-associated proteins are shown in grey. Circle diameter indicates relative protein size. Blue: 50S ribosomal proteins; yellow: 30S ribosomal proteins; green: RNAP; orange: NusA. Each edge represents one or more crosslinks. (**B**) Network showing interactors of RNAP and NusA. The NusA N-terminal domain, S1 domain, RNA binding KH domains, proline rich region (PR) are annotated. Line thickness represents the number of identified crosslinks.

The *M. pneumoniae* RNAP core consisting of the conserved subunits α, β and β’, was found to interact with the known auxiliary factors SigA, GreA, NusG, NusA, SpxA, and RpoE (firmicute-specific RNAP δ subunit) (*19*) (Fig. 1B). Additionally, two uncharacterized, essential proteins MPN555 and MPN530 were found (*20*). MPN555 crosslinked to the C-terminal region of the β subunit, and was recently shown to have a DNA binding profile similar to RNAP in ChIP-seq (*16*). We further validated the interaction between MPN530 and β/β’ subunits by a bacterial two-hybrid screen (fig. S4). Despite these interactions, no crosslinks between the RNAP core and the ribosome were identified, which would have been expected if a putative transcription-translation coupling supercomplex in *M. pneumoniae* had a similar architecture to either of the previous *in vitro* structures (*6, 7*). Interaction between NusG and the ribosomal protein S10, previously reported in *E. coli*, was also not detected (*8, 9*).

Intriguingly, NusA, a transcription elongation factor involved not only in elongation, but also termination and antitermination (*21, 22*), was found to interact with RNAP via its N-terminus, and with the mRNA entry site of the ribosome via its C-terminal region (Fig. 1B). Our data indicated a potential architecture in which RNAP and the ribosome are linked by NusA.

To investigate the 3D structure of this potential indirect RNAP-ribosome association, cryo-ET data were acquired on unperturbed, frozen-hydrated *M. pneumoniae* cells (Fig. 2A and fig. S5A-B). Ribosomes were localised by template matching with a *de novo*-generated ribosome reference (fig. S5D-F). 108,501 sub-tomograms were extracted from 352 tomograms and subjected to averaging and classification (fig. S6). As expected, ribosomes exhibited large structural heterogeneity and were first sorted into classes representing the 50S/large ribosomal subunit (30.3%), 70S ribosomes (53.3%) and 70S ribosomes closely assembled in a polysome configuration (16.4%) (Fig. 2A and fig. S7). Subjecting 73,858 sub-tomograms of all 70S ribosomes to a new sub-tomogram analysis workflow enabling 3D average-based tilt-series alignment (fig. S6) resulted in a 5.6 Å ribosome density (Fig. 2B, fig. S8 and supplementary text). We fitted the obtained ribosome density with a homology model based on the *Bacillus subtilis* ribosome (PDB 3J9W) which enabled mapping of most *M. pneumoniae* ribosomal proteins (19 30S/small subunit and 28 50S/large subunit r-proteins) (fig. S9). Unaccounted densities at the C-termini of the 50S r-proteins L22 and L29 (Fig. 2B, insert) were assigned to two α-helices from C-terminal extensions that are unique to *M. pneumoniae* and its close relatives (fig. S10 and S11). A third protein, L23, also containing a C-terminal extension in *M. pneumoniae* was predicted to be unstructured and did not produce a discernible density in the map (fig. S12). To the best of our knowledge, the herein reported resolution is unprecedented in cellular cryo-ET of asymmetric structures, enabling, in conjunction with CLMS, *de novo* determination and assignment of secondary structure elements *in cellulo* (fig. S9-12).

**Fig. 2.**
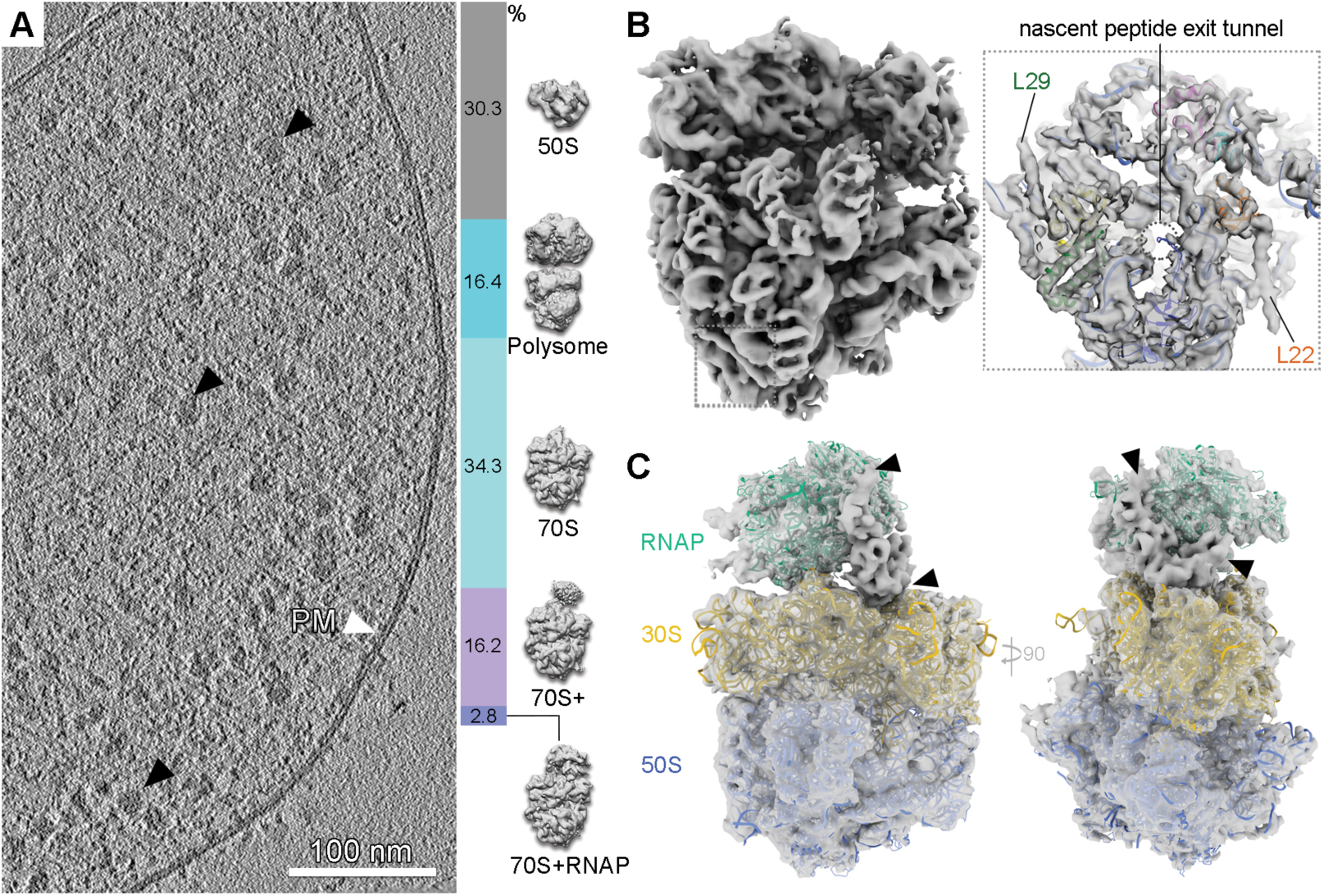
In-cell ribosome structures determined by cryo-ET and sub-tomogram analysis reveal the presence of a RNAP-ribosome supercomplex. (**A**) Tomographic slice of a *M. pneumoniae* cell. Black arrowheads: ribosomes. PM: plasma membrane. Right: classification of 108,501 ribosome sub-tomograms from intact *M. pneumoniae* tomograms. (**B**) Left: 5.6 Å in-cell 70S ribosome density; insert: density near the peptide exit tunnel (dashed circle) shows two helices not accounted for in the fitted homology model (L22 and L29). (**C**) 9.2 Å in-cell density obtained from the class representing RNAP-ribosome supercomplexes (2.8% in (A)). Homology models are fitted and the unassigned density indicated (black arrowheads).

Focused classification of the 70S class near the mRNA entry site detected a class of ribosomes with additional density (70S+, Fig. 2A). A second class contained a density resembling the known RNAP structure (70S+RNAP, Fig. 2A and fig. S7). Refinement of this class provided a 9.2 Å density (fig. S13), into which the ribosome and RNAP models could be unambiguously fitted (Fig. 2C). In line with the in-cell CLMS data, this map contained unexplained density at the interface between the two complexes (Fig. 2C, arrowheads). Multi-body refinement allowed to better resolve the RNAP density and revealed a large degree of conformational flexibility (fig. S14 and Movie 1). Nevertheless, the path of DNA and about one turn of the DNA-RNA hybrid duplex were clearly visible, demonstrating that RNAP was in an elongating state (Fig. 3A and fig. S15C). No density was observed for single-stranded or folded mRNA near the exit tunnel. This, together with the presence of the 70S ribosome, indicated that the supercomplex represents an actively elongating expressome.

**Fig. 3.**
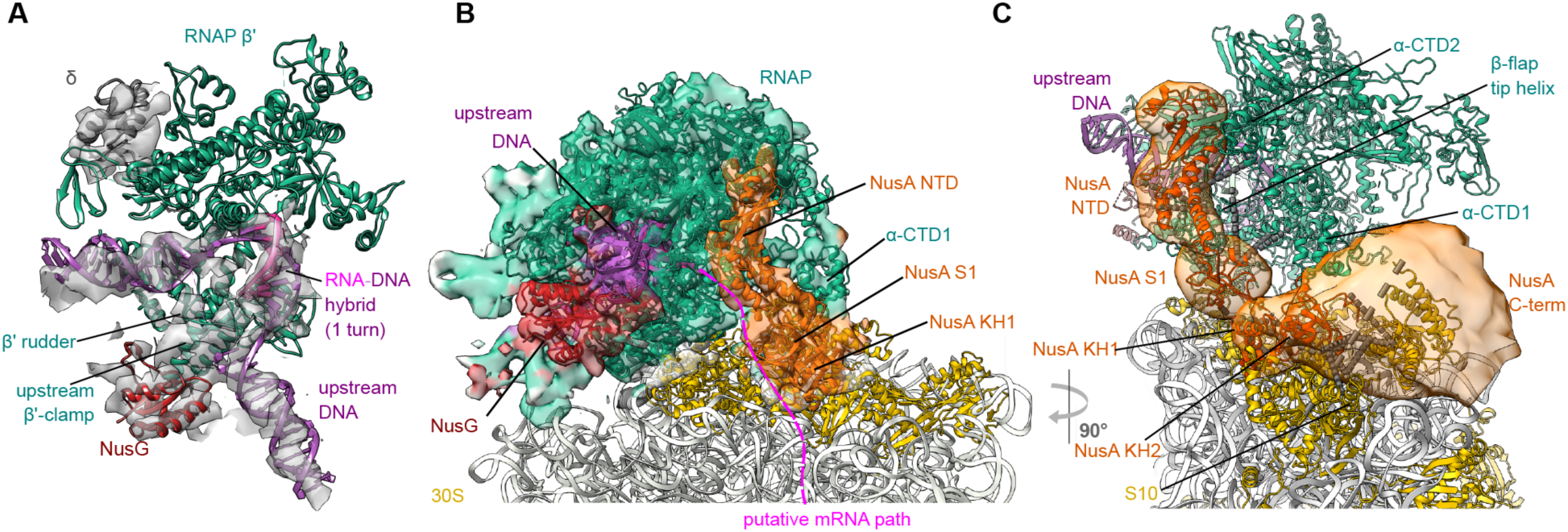
Integrative model of the elongating expressome in *M. pneumoniae*. (**A**) The density in the RNAP region of the multibody-refined expressome map corresponding to the DNA, RNA-DNA hybrid and upstream β’ clamp, δ subunit and NusG are shown. It accommodates one turn of the RNA-DNA hybrid (PDB 6FLQ), consistent with an elongating polymerase. (**B**) Integrative model with cryo-EM density of the RNAP-NusG-NusA-ribosome elongating expressome. Structured regions modeled as rigid bodies are represented as coloured cartoons. Coarse-grained regions are not shown. The density colors represent the subunits occupying the corresponding volumes in the selected model. Schematic of the putative mRNA path refers to the shortest distance between mRNA exit and entry sites. (**C**) Back-view of the mRNA exit tunnel face of RNAP covered by NusA. Inter-protein crosslinks involving NusA are displayed as grey dashed lines. The localisation probabilities for the NusA domains and C-terminal region are displayed as orange densities. The NusA NTD-S1 domains localise near the mRNA exit tunnel of RNAP, KH domains sit on top of mRNA entry channel of the ribosome and the C-terminal region is localised over the face of the 30S subunit.

Both CLMS and cryo-EM data showed binding of the elongating factor NusG to its conserved site near the upstream DNA and the β’ clamp helices of RNAP (fig. S15, S16) (*23*). *M. pneumoniae* NusG contains large inserts of unknown structure in its CTD, but does retain the residues involved in the NusG-S10 (NusE) interaction (*8*), suggesting that this interaction may be conserved in *M. pneumoniae*. However, the arrangement of RNAP relative to the ribosome places NusG away from S10, indicating that this interaction does not occur in the elongating expressome (fig. S16). The CLMS data indicate that SigA, SpxA and GreA bind to their canonical sites on RNAP, but these proteins did not produce detectable density in the elongating expressome map (fig. S17), consistent with them being part of other RNAP complexes.

The remaining unaccounted density between RNAP and the ribosome is consistent with NusA binding to RNAP via its N-terminal domain, as suggested by CLMS (Fig. 1B and fig. S18). To better understand the arrangement of the NusA domains and flexible regions of RNAP, CLMS data and the cryo-EM density were used to derive a model of the elongating expressome using the Integrative Modeling Platform (*24*) (fig. S19). Each domain of NusA, the RNAP core, the β flat-tip helix and δ subunits, were coarse-grained in a multi-scale representation as separate rigid bodies (table S4 and S5). *M. pneumoniae* NusA contains a C-terminal extension not found in *E. coli* or *B. subtilis*, which is proline-rich and predicted to be disordered (fig. S18 and supplementary text). This region, found to be crosslinked to multiple 30S r-proteins including S10, was left flexible in the modeling and not fitted into the cryo-EM density. The structured regions of NusG, the RNAP core and the ribosomal proteins were unambiguously fitted into the density as rigid bodies prior to modeling of the flexible regions (fig. S19).

The best-scoring solutions (fig. S20) showed an architecture in which the N-terminal and S1 domains of NusA bind RNAP similarly to the *E. coli* paused elongation complex (*22*), with the S1 domain near the RNAP mRNA exit tunnel (Fig. 3B, C and fig. S21). The two KH domains were positioned near S3, S4 and S5 at the mRNA entry channel of the 30S subunit. The orientation of KH domains keeps their RNA-binding interface in a position that could interact with the nascent mRNA, in a binding mode similar to previously reported structures (*25*) (fig. S22). The CTDs of α subunits were localised between the NusA NTD and the second KH domain, with one α-CTD fitting a region of density between the second NusA KH domain and the RNAP core (fig. S22 and supplementary text). Due to their flexible nature, the α-CTDs were found to arrange in a wide range of conformations. Additionally, the regions of RNAP for which no template structure was available fit into the unaccounted density near the downstream DNA. The loop comprising residues 48-91 of NusG localises in the density below the DNA entry channel. The RNAP δ subunit is positioned according to crosslink information in the density below RNAP β’ CTD (fig. S23), which is consistent with its suggested role in regulating RNAP-DNA interaction (*19*).

The integrative model suggests that NusA forms a bridge between elongating RNAP and ribosome in the active expressome. To determine whether this architecture requires active translation elongation, we collected cryo-ET data on cells treated with chloramphenicol (Cm). The percentage of 70S ribosomes increased dramatically compared to untreated cells (Fig. 4A and fig. S24), while the percentage of the polysome class remained similar. The resulting 6.5 Å Cm-treated ribosome density (Fig. 4B) had well-resolved tRNAs in both A and P sites, in agreement with a previous Cm-stalled ribosome structure (PDB 6ND5; Fig. 4C) (*26*). Further sub-tomogram analysis did not detect RNAP density near the 30S mRNA entry site, indicating that stalling ribosomes leads to dissociation of the active expressome.

**Fig. 4.**
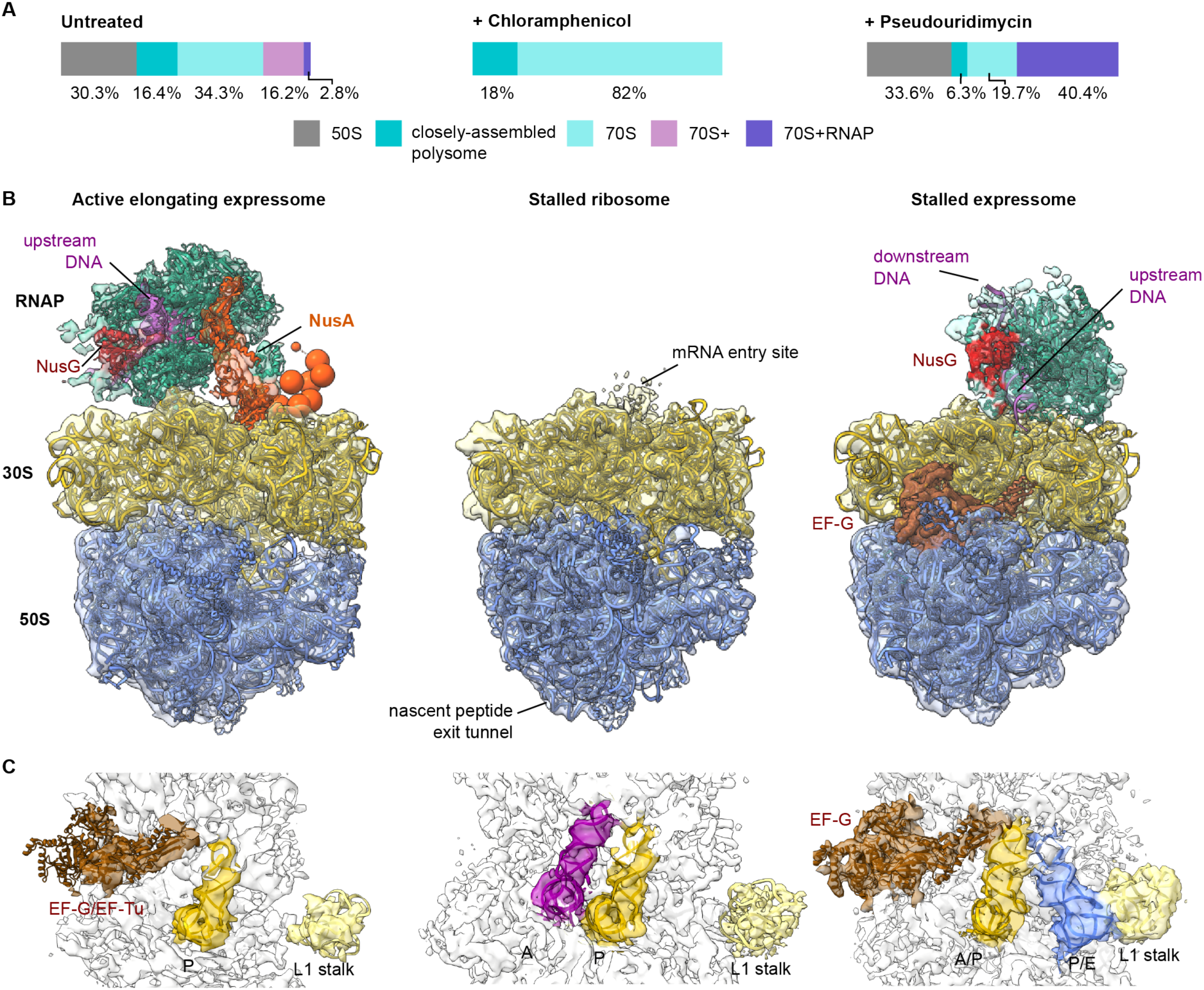
Stalling translation or transcription alter the expressome architecture in cells. (**A**) Classification of sub-tomograms in untreated, Cm-treated, and PUM-treated *M. pneumoniae* cells reveal large shifts in ribosome populations following perturbations. (**B**) Models of RNAP-ribosome supercomplexes. Left: In untreated cells, the ribosome is captured in its actively translating state. The density is fitted with the integrative model of the elongating expressome. Center: Cm decouples the ribosome and RNAP. Right: in PUM-treated cells, the ribosome encounters the stalled RNAP and is stalled in the pre-translocation state. (**C**) tRNA occupancies in densities derived from untreated, Cm-treated, and PUM-treated cells. In untreated cells, densities for P-site tRNA and elongation factors are visible. Upon addition of Cm, A and P-site tRNAs are observed. In PUM-treated cells, presence of EF-G and hybrid A/P* and P/E site tRNA suggest the ribosome is stalled in the pre-translocation state.

The dependence of expressome integrity on active transcription was probed by treating cells with the specific RNAP inhibitor pseudouridimycin (PUM) (*27*), which significantly increased the percentage of well-resolved expressome complexes from 2.8% in untreated cells to 40% (Fig. 4A). The PUM-induced expressome was refined to 7.1 Å (fig. S25 and S26), revealing direct interaction between RNAP and the ribosome, and excluding any density for NusA. The δ subunit and NusG remained bound to RNAP (Fig. 4 and fig. S27C, D). Interestingly, tRNAs in the ribosome were found in hybrid A/P* and P/E states, and density corresponding to EF-G was well-resolved (Fig. 4B). These results suggest that ribosomes are trapped in a pre-translocation state (*28*) unable to complete the translocation cycle, presumably due to the stalled RNAP. This stalled expressome closely resembles the architecture of the *E. coli* expressome solved *in vitro* (*6*) (fig. S27). Taken together, these results indicate that active transcription and translation elongation are required for the native RNAP-NusG-NusA-ribosome elongating expressome architecture.

The in-cell architecture of the active expressome derived here elucidates how physically coupled transcription and translation elongation are structurally arranged in *M. pneumoniae*. Our data also highlights the structural heterogeneity of the process inside cells (fig. S7, Movie 1), consistent with the presence of multiple, context-dependent coupling mechanisms. We mapped both NusG and NusA, two general and conserved factors that can bind to RNAP throughout transcription elongation (*10*). In this expressome architecture, where both RNAP and the ribosome are elongating, NusG cannot interact with the ribosomal protein S10. Instead, NusA is found at the RNAP-ribosome interface following the putative nascent mRNA path. There NusA may act as a sensor for a translating ribosome approaching RNAP, and could modulate transcription elongation to achieve transcription-translation coupling. This model is consistent with the described versatile roles of NusA in directing mRNA during pausing, termination and antitermination (*1, 22, 29*). These findings highlight the enormous potential of integrative in-cell structural biology approaches in elucidating dynamic and complex cellular processes within their native context.

## Acknowledgments

We are grateful to Mahmoud Chaabou, Helena Barysz, Zhuo Angel Chen, Lutz Fischer, Colin Combe, Martin Graham, and Ievgeniia Zagoriy. Jan Kosinski, Christoph Müller and Martin Beck are acknowledged for critical reading of the manuscript.

## Author contributions

F.O., L.X., J.M. and J.R. designed the experiments; F.O., L.S. and S.L. collected and processed CLMS data; C.B. and J.S. performed the two-hybrid work; L.X. and J.M. collected and processed cryo-EM data; W.H. supported cryo-EM data collection; D.T. and P.C. contributed new cryo-EM processing software; A.G. and S.L performed integrative modelling; F.O., L.X., A.G., J.M. and J.R. prepared figures and wrote the manuscript with input from all authors.

## Data and materials availability

The Supplementary Materials contain additional data. EM densities have been deposited in the EMDataBank with the following accession numbers: EMD-10677:10687. CLMS data are available via ProteomeXchange with identifiers PXD017711 (DSSO) and PXD017695 (DSS). Integrative modeling results are uploaded to PDB-dev.

## Supplementary Materials

Materials and Methods

Figures S1-S27

Tables S4-S6

References (*30-91*)

## Other Supplementary Materials

Movies S1-S2

External Data Tables S1-S3

