## Supplementary Material for "In-cell architecture of an actively transcribing-translating expressome"

### Materials and Methods

*Mycoplasma pneumoniae* cultivation for CLMS

*M. pneumoniae* strain M129 (ATCC 29342) in the 32nd broth passage was used. Bacteria were grown at 37°C in 150 $cm^2$ tissue culture flasks containing 80 ml of modified Hayflick medium (*30*): 14.7 g/L Difco™ PPLO (Becton Dickinson, USA), 20% (v/v) Gibco™ horse serum (New Zealand origin, Life Technologies, Carlsbad, USA), 100 mM HEPES-Na (4-(2-hydroxyethyl)-1-piperazineethanesulfonic acid; pH 7.4), 1% (w/w) glucose as a carbon source, 0.002% (w/w) phenol red and 1,000 U/mL penicillin G.

Crosslinking of the Cells

Cells were harvested from media by centrifugation at 16,000 g for 5 min, followed by two washes with 500 µl 1x Phosphate-buffered saline (PBS) buffer. Crosslinker was dissolved freshly in water-free Dimethyl sulfoxide (DMSO) at 50 mM. The crosslinker, either disuccinimidyl suberate (DSS, Thermo Scientific) or disuccinimidyl sulfoxide (DSSO, Thermo Scientific) was quickly added to cells resuspended to a final concentration of 1 mg/ml cells in 1x PBS at a final concentration of 2 mM (5% DMSO in final volume). The cells were incubated at 25°C for 30 min under gentle agitation, followed by quenching with 50 mM Tris-HCl (pH 7.5) for 15 min. Cells were pelleted and snap-frozen with liquid $N_2$ before storing at -80°C.

Cell lysis and Proteolysis for LC-MS Analysis

For proteome extraction, the cell pellet was lysed at 20 mg/ml in denaturing solution (6 M Urea, 2 M Thiourea, 50 mM ammonium bicarbonate (ABC, Sigma), 1 mM dithiothreithol (DTT, Sigma)), aided by sonication using a 3.2 mm Qsonica microtip probe (output 1.5, duty cycle 0.2; 20 min per sample; on ice).

Protein concentration of the cell lysate was assessed by Bicinchoninic acid assay (BCA, Pierce) according to the manufacturer's instructions and was diluted to 1 mg/ml with the denaturing solution. The lysate was reduced for 30 min by adding 10 mM DTT, followed by addition of 25 mM iodoacetamide (Sigma) and incubation in the dark for 20 min. The reaction was quenched by adding 10 mM DTT. Proteolysis was carried out using LysC (Pierce) at 1:100 (mass ratio, protease:protein) for 4 h at 37 °C, followed by 1:5 dilution with 50 mM ABC and tryptic digestion (1:50 (m/m), Pierce) for 16-18 h at 37°C. The reaction was stopped by adding 10% trifluoroacetic acid (TFA, Honeywell) to pH 2-3. Samples were cleaned-up via Stage-tipping (3M) and stored at -20°C until further use, i.e. peptide fractionation.

Peptide fractionation by Strong Cation Exchange Chromatography (SCX)

Peptide digests were eluted from Stage-tips, dried using a vacuum concentrator and resuspended in SCX running buffer A (30% acetonitrile (ACN, VWR Gmbh), 10 mM $KH_2PO_4$ pH 3.0 (Sigma)). Separation of peptides was accomplished using a non-linear gradient with running buffer B (30% ACN, 1 M KCl, 10 mM $KH_2PO_4$, pH 3.0), as described (*31*). For the fractionation, a Shimadzu HPLC system (LC-20AD, SIL-20A HT, SPD-M20A, CBM-20A) connected to a PolyLC Polysulfoethyl A column (100 x 2.1 mm, 3 µm, 300 Å) was used. Fractions of 200 µl each were collected over the elution window (ca. 18 column volumes). Collected fractions of interest from five runs were pooled, desalted using Stage-Tips and stored at -20°C.

Peptide fractionation by Size-Exclusion Chromatography (SEC)

Dried peptide digests or fractions were resuspended in SEC running buffer (30% ACN, 0.1% TFA) and injected onto a Superdex Peptide 3.2/300 column running at 50 µl/min flow rate connected to an Äkta Pure system. Fractions of 100 µl were collected and dried in a vacuum concentrator (*17*, *32*) and stored at -20°C.

Peptide fractionation by hydrophilic Strong Anionic Exchange Chromatography (hSAX)

Peptides from Stage-Tips were dried and subsequently resuspended in hSAX running buffer A (20 mM Tris-HCl, pH 8.0). Sample was injected onto a Dionex IonPac AS24 column (2 x 250 mm) running with 150 µl/min operated at 15°C under Unicorn 7.1 on an Äkta Pure system. Elution of peptide mixtures was achieved by applying an exponential gradient with buffer B (20 mM Tris-HCl, pH 8.0, 1 M NaCl; gradient identical to SCX gradient), while collecting fractions of 150 µl. Obtained fractions of interest were acidified with TFA to pH 2-3, pooled, desalted by Stage-tipping and stored at -20°C.

Two-dimensional peptide fractionation for each CLMS dataset

Four datasets were collected to achieve significant depth of analysis (Fig. S1A).

Dataset 1 - Peptides from cells crosslinked with DSS were fractionated by SCX. Then 11 fractions enriched for crosslinked peptides were fractionated individually by SEC as a second dimension. Three fractions enriched for crosslinked peptides from each SEC run were collected, resulting in 33 total fractions for MS acquisition.

Dataset 2 - Proteins were extracted and subjected to native SEC, as described in (*33*), and all fractions with protein complexes of molecular weight larger than 150 kDa were pooled for digestion and peptide separation as in Dataset 1.

Dataset 3 - Peptides from cells crosslinked with DSS were fractionated by SCX. Then 9 fractions enriched for crosslinked peptides were fractionated individually by hSAX into 8 pools as a second dimension. This resulted in 72 total fractions for MS acquisition.

Dataset 4 - Peptides from cells crosslinked with DSSO were fractionated by SCX to give 9 fractions enriched for crosslinked peptides. Each fraction was fractionated individually by hSAX, resulting in 9x15 total hSAX fractions.

LC-MS/MS acquisition of crosslinked samples

Acquisition of crosslinked peptide spectra was performed on a Fusion Lumos Tribrid Mass Spectrometer (Thermo Fisher Scientific, San Jose, USA) connected to an Ultimate 3000 UHPLC system (Dionex, Thermo Fisher Scientific, Germany). Mobile phase A (0.1% (v/v) formic acid), and mobile phase B (80% (v/v) acetonitrile, 0.1% (v/v) formic acid). The samples were dissolved in 1.6% acetonitrile (Fluka), 0.1% formic acid (Fluka) and separated on an Easy-Spray column (C-18, 50 cm, 75 μm internal diameter, 2 μm particle size, 100 Å pore size) running with 300 nl/min flow rate using optimized gradients for each offline-fraction (ranging from 2% B to 55% B over 62.5, 92.5 or 152.5 min (for datasets 1, 2-3, and 4 respectively), then to 55% in 2.5 min and to 95% in 2.5 min). Fractions were injected twice when more than 3 μg peptide material was available. The MS data were acquired in data-dependent mode using the top-speed setting with a 3 second cycle time. For every cycle, the full scan mass spectrum was recorded in the Orbitrap at a resolution of 120,000 in the range of 400 to 1,600 m/z. Ions with a precursor charge state between +3 and +6 were isolated and fragmented. The fragmentation regime depended on the crosslinking reagent employed: For the non-cleavable linker DSS, Higher-energy collisional dissociation (HCD) energies optimized for mass and charge of a precursor species were applied (*34*). For the cleavable DSSO crosslinker, stepped fragmentation energies of 18, 24 and 30 normalized collision energy (NCE) were used that were found appropriate (*35*). The fragmentation spectra were then recorded in the Orbitrap with a resolution of 30,000. Dynamic exclusion was enabled with single repeat count and 60 second exclusion duration.

Identification and statistical validation of crosslinked peptides

Raw data from mass spectrometry was processed using MSconvert (version 3.0.11729) (*36*) including denoising (top 20 peaks in 100 m/z bins) and conversion to mgf-file format. Masses of precursor peptides were recalibrated. Obtained peak files were analysed using xiSearch 1.6.746

(*17*) with the following settings: MS1/MS2 error tolerances 4 and 10 ppm, allowing up to 3 missing isotope peaks (Lenz et al., 2018), tryptic digestion specificity, carbamidomethylation on Cys as fixed and oxidation on Met as variable modification, losses: –$CH_3SOH$ / –$H_2O$ / –$NH_3$, crosslinker DSS (138.06807 Da linkage mass) or DSSO (158.0037648 Da linkage mass) with variable crosslinker modifications ("DSS-NH2" 155.09463 Da, "DSS-OH" 156.07864 Da, "DSSO-NH2" 175.03031 Da, "DSSO-OH" 176.01433 Da). For samples crosslinked with DSSO, additional loss masses were defined accounting for its cleavability ("A" 54.01056 Da, "S" 103.99320 Da, "T" 85.98264). Crosslink sites for both reagents were allowed for side chains of Lys, Tyr, Ser, Thr and the protein N-terminus. The database consisted of 687 Swiss-Prot annotated entries for *M. pneumoniae* strain ATCC 29342/M129 (taxon identifier 272634), 19 small open reading frames that were recently annotated (*20*), and annotated lysine acetylation sites (*37*). Decoy sequences were generated by reversing the protein sequences.

Results were filtered to an estimated false-discovery rate (FDR) of 5% on residue-pair-level and 5% on protein-protein interaction (PPI)-level using xiFDR (version 1.2.30.59) (*18*). Inter- and intra-crosslinks were handled separately for FDR estimation. The DSS datasets (datasets 1-3) were validated together for FDR estimation. Spectral matches in the DSSO dataset were prefiltered before FDR estimation to only those that had cleaved crosslinker peptide fragments for both peptides.

<u>LC-MS/MS acquisition and data analysis for bottom-up proteomics</u>

For identifying proteome abundances we subjected the proteome to bottom-up proteomics. The Proteome was extracted as outlined above. Mass spectrometric analysis of native (non-crosslinked) *M. pneumoniae* proteome was performed on a Lumos Tribrid Mass Spectrometer (Thermo Fisher Scientific, San Jose, USA) connected to an Ultimate 3000 RSLCnano system (Dionex, Thermo Fisher Scientific, Germany). The proteome was analysed in triplicate, each with a total run time of 230 min. The chromatographic setup was identical to the one used for crosslinking acquisitions with the following changes on the LC gradient: The gradient started at 2% B to 5% B in 1 min, to 7.5% B in 5 min, then to 32.5% in 190 min, 40% B in 7 min, 50% B in 2.5 min followed by ramping to 90% B in 2.5 min and washing for 5 min. The settings of the mass spectrometer were as follows: 2 second cycle time; Data-dependent mode; MS1 scan at 120,000 resolution over 350 to 1,600 m/z; automatic gain control (AGC) target of $3e^6$ with maximum Injection time (IT) of 50 ms; MS2 triggered only on precursors with charge state between +2 and +7; 1.6 m/z isolation width; AGC target of $1e^5$ with 80 ms max. IT and min. AGC target of $2.5e^4$; fragmentation by HCD 29; MS2 scan was ion trap in Rapid mode; peptide match was set as preferred and dynamic exclusion was enabled upon single observation for 60 seconds.

Mass spectrometry raw data was processed using MaxQuant 1.6.1. with default settings with minor changes: three allowed missed cleavages; up to four variable modifications per peptide including oxidation on Met, acetylation on protein N-terminal. Carbamidomethylation on Cys was set as fixed. The database covered 706 proteins (identical to crosslinking search, see above). The 'matching between runs' feature was disabled with default settings. Protein quantification was based on two or more peptides using the iBAQ approach (*38*). Proteins are reported at 1% FDR (Table S2).

<u>*Bacterial two-hybrid assay (BACTH)*</u>

Primary protein-interactions were validated by bacterial two-hybrid analysis (*39*). The BACTH system is based on the interaction-mediated reconstruction of Bordetella pertussis adenylate cyclase (CyaA) activity in *E. coli*. Functional complementation between two fragments (T18 and T25) of CyaA as a consequence of the interaction between bait and prey proteins results in cAMP synthesis. The β-galactosidase reporter gene was fused to a cAMP-dependent promoter which allows indirect detection of the protein-protein interaction by assaying β-galactosidase activity. Plasmids pUT18/pUT18C and p25N/p25 allow the expression of proteins fused to the C- or N-terminal T18 and T25 fragments of CyaA, respectively. For these experiments, we constructed the plasmids pGP3280-3297, pGP295-300, pGP1451-GP1453. DNA fragments corresponding to the genes mpn530, sigA (mpn352), rpoA (mpn191), rpoB (mpn516), rpoC (mpn515), and spxA (mpn266) were obtained by PCR, digested and ligated into the vectors pUT18/pUT18C and p25N/p25 (Table S3). These plasmids were used for co-transformation of *E. coli* BTH101, and the protein-protein interactions were then analyzed by plating the cells on LB plates containing 100 µg/ml ampicillin, 50 µg/ml kanamycin, 80 µg/ml X-Gal (5-bromo-4-chloro-3-indolyl-β-D-galacto-pyranoside), and 1 mM Isopropyl β- d-1-thiogalactopyranosideIsopropyl β-d-1-thiogalactopyranoside (IPTG). The plates were incubated for a maximum of 36 h at 28°C.

<u>*M. pneumoniae* cultivation and vitrification for cryo-electron microscopy (EM)</u>

Wild-type *M. pneumoniae* strain M129 (ATCC 29342） were cultivated in cell culture flasks (Greiner Bio-One, Frickenhausen, Germany) with modified Hayflick medium (*30*) as described above. For cryo-EM grid preparation, 200 mesh gold (Au) grids with holey Quantifoil support (R2/1, Quantifoil Micro Tools, Jena, Germany) were glow discharged for 45 s and sterilized under UV irradiation in a laminar flow hood for 30 min, before immersing into culture medium. After inoculation, *M. pneumoniae* cells were grown at 37°C for less than 20 hours to ensure that cells were in a fast-growing phase. Vitrification was carried out using a manual plunger (Manufactured by the Max Planck Institute of Biochemistry, Martinsried, Germany). Grids with cells were quickly washed with PBS solution containing 10 nm protein A-conjugated gold beads (Aurion, Wageningen, Netherlands), blotted for 2 seconds from the back side, and plunged into a liquid ethane/propane mixture at liquid $N_2$ temperature. For antibiotic-treated datasets, either chloramphenicol (Cm; Sigma-Aldrich, USA) at a final concentration of 0.5 mg/ml or pseudouridimycin (PUM; Adipogen AG, Switzerland) at a final concentration of 0.4 mg/ml were added directly into the culture medium 15-20 minutes prior to vitrification. The frozen grids were stored in sealed boxes in liquid $N_2$ until further processing.

Cryo-electron tomography

Cryo-electron tomography data were collected on a Titan Krios microscope operated at 300 kV (ThermoFisher Scientific, Eindhoven, Netherlands) equipped with a field-emission gun, a Quantum post-column energy filter (Gatan, Pleasanton, CA, USA), a K2 Summit direct detector camera (Gatan) and a Volta phase plate (VPP). Data were recorded in dose-fractionation mode using acquisition procedures in SerialEM software v3.7.2 (*40*). Prior to tilt-series acquisition, montages of grid squares were acquired at 2.2 nm/pixel (Fig. S5A). Tilt-series were collected with a dose-symmetric scheme (*41*), Energy-Filtered TEM operated in zeroloss, nano-probe mode, magnification 81,000x, calibrated pixel size at the specimen level of 1.7 Å, defocus range 1.5 to 3.25 µm, tilt increment 3° with constant dose for all tilts, tilt range -60° to 60°, total dose of ~ 120 $e^{-}/Å^{2}$. 14 tilt-series were acquired with the Volta phase plate (VPP) (Fig. 2A), primarily for generation of a *de novo* reference for template matching (described below). Alignment and operation of the VPP were essentially carried out as described previously, applying a beam tilt of 10 mrad for autofocusing (*42*). Tilt-series for high-resolution sub-tomogram analysis were acquired without VPP, with a 70 µm objective aperture and a beam tilt of 4 mrad for autofocusing (Fig. S5B). In total, 500 tomograms were acquired: 352 for native, untreated cells, 65 for Cm-treated and 83 for PUM-treated cells.

Tomogram reconstruction

Prior to tilt-series alignment, movies from the K2 camera were corrected for beam-induced motion with the SerialEM plugin. Tilt-series alignment using fiducial gold beads and tomographic reconstruction were performed in etomo from the IMOD software package v4.9 (*43*). Aligned images were 4 times binned to a pixel size of 6.8 Å. For tomographic reconstruction, the radial filter options were left as default (cut off, 0.35; fall off, 0.05). Cell thickness was estimated using central YZ-slices of the tomograms. The average cell thickness estimated from the tomograms was 158 nm (Fig. S5C).

Ribosome localization using template matching

To generate a data-driven template for correlation-based ribosome localization as implemented in pyTOM (*44*), 404 ribosomes were manually selected from 7 VPP tomograms using EMAN2 boxer (*45*). 4x binned sub-tomograms were extracted and refined in RELION (version 3.0.7) using a featureless sphere as an initial reference. The refinement converged into a ribosome density (Fig. S5D). The density was low-pass filtered to 30 Å resolution using TOM toolbox (*46*) implemented in MATLAB (MathWorks 2016), and used for template matching in 4x binned conventional defocus tomograms. For each tomogram, the 400 highest-scoring cross-correlation peaks were extracted, with a minimal Euclidean distance of 20.4 nm (30 voxels) between peaks to avoid multiple detection events for the same ribosome. The peaks were visually inspected to exclude obvious false positives, such as membranes and edges of the carbon support. With an initial dataset of 20 defocus tomograms and 4,668 sub-tomograms, a ribosome density of 25 Å resolution was obtained. The ribosome density was low-pass filtered to 30 Å and used as the new template for matching in all defocus tomograms. For each tomogram, the top 400 hits were visually inspected, and on average 307 hits were retained after removing obvious false positives (Fig. S5E, F). In total, 108,501 ribosome sub-tomograms were extracted from the untreated cells tomograms, 21,299 from Cm-treated and 22,089 from PUM-treated datasets.

Sub-tomogram analysis

The sub-tomogram analysis workflow described below is summarized in Figure S6. All analysis steps were performed using Warp (version 1.0.6) and RELION (version 3), mostly following previously published protocols (*47*, *48*).

Defocus parameters were determined using Warp on individual tilt images and consecutively refined on the whole tilt-series following tilt-series alignment in etomo. Phase modulations introduced by the contrast transfer function (CTF) and radiation damage were corrected using 3D CTF/wedge models reconstructed for each sub-tomogram in Warp. Ribosome sub-tomograms based on coordinates retrieved by pyTOM template matching were reconstructed at 2x binning (3.4 Å/pix) in Warp. These were subsequently sorted into different classes using RELION 3D classification (*49*). Sub-tomograms that could not be aligned and 50S ribosomes were excluded first, and 70S ribosomes were further classified to separate those arranged in a closely assembled polysome configuration (Fig. S7). Subsequent rounds of focused 3D classification with a mask at the 30S mRNA entry site enabled classifying the remaining 70S sub-tomograms into 3 major subclasses: ribosomes with well-defined RNAP density near the mRNA entry site, ribosomes with undefined additional density, and ribosomes without additional density (Fig. S7). At least 3 independent classification runs with different masks and RELION settings were performed to validate the convergence of the classification results. Refinement using RELION 3D auto-refine focused on the ribosome density of 73,858 70S ribosomes attained a density at 10.3 Å resolution as determined by gold standard Fourier shell correlation (FSC) at 0.143. To improve the resolution, the sub-tomogram poses and tilt-series geometry were refined in a new tool associated with Warp, *M* (see details in the following section). With the same sub-tomogram dataset re-extracted at the original pixel size of 1.7 Å after *M* refinement, the final global resolution improved to 5.6 Å (Fig. S8A, B). Local resolution map, angular distribution analysis, as well as 3D FSC analysis (*50*), validated the reported isotropic resolution (Fig. S8 C-E), while densities corresponding to secondary structural elements confirmed the quality of the sub-tomogram average (Fig. S8 F-H).

Different refinement strategies were performed with the 2,952 ribosomes exhibiting clear RNAP density. Refinement with a spherical mask generated a density of 8.5 Å global resolution (Fig. S13A). However, the resolution of the RNAP density was in the range of ~ 12 Å (Fig. S13B). To avoid influence of the ratcheting movement between the small and large ribosomal subunits, while keeping the RNAP-30S interface fully covered, a focused refinement with a local RNAP-30S mask was carried out (Fig. S13C). Although the global resolution dropped to 9.2 Å, the density quality of the RNAP was improved (Fig. S9D). High flexibility of the RNAP itself and the interface with the small ribosomal subunit hindered achieving higher local resolution. To overcome this, multi-body refinement as implemented in RELION was performed (*51*). Following one consensus refinement, 3 bodies corresponding to the three major parts of the supercomplex were segmented (i.e. RNAP and linker density, small and large ribosomal subunits), and a local mask was created for each body (Fig. S14A-D). Principal component analysis (PCA) on the relative orientations of all bodies was performed to characterise the most predominant motions (Fig. S14E and Movie 1).

Similar sub-tomogram analyses, including classification, refinement and multi-body refinement were performed for the Cm-treated dataset (Fig. S24) and PUM-treated dataset (Figs. S25, S26 and Movie 2). The resolution of all sub-tomogram averages was estimated based on FSC of two independent halves of the data using FSC=0.143 as the cutoff criterion.

Warp and M processing pipeline for sub-tomogram analysis

To obtain higher resolution for the 3D averages, the particle (sub-tomogram) poses and tilt-series alignments were refined in a new software tool, *M*. Rather than following single particle analysis' central paradigm of treating each particle as a single, isolated entity, *M* implements a multi-particle analysis approach. All particles in a single tilt-series are treated as parts of the same physical system, which has its own set of hyperparameters that are optimized simultaneously with the individual particle poses.

In the case of tilt-series, the hyperparameters include the global rotation angles, coarse non-linear image-space deformation to describe stage movement and beam induced motion, and the CTF models of individual tilt images, as well as a non-linear 3D deformation model of the tomographic volume describing its local changes as a function of the accumulated dose with a fine spatial, and coarse temporal resolution. Making use of the high amount of signal available for each ribosome particle, the particle poses themselves were modeled as a function of accumulated dose, with 3 optimizable sets of rotation and translation parameters per particle.

The non-linear deformation models were implemented as uniform 2D (image deformation in each tilt) and 4D (volume deformation over dose) grids of cubic splines, as described previously (*48*). The coarseness of the deformation models had a regularizing effect on the local divergence of individual 2D particle tilt image rotations and shifts, as previously formulated (*52*), and implemented in emClarity (*53*) and, for the special case of 2D data, Bayesian particle polishing in RELION 3.0 (*54*). Multi-particle refinement in *M* provides the first unified framework for performing reference-based particle alignment with physically plausible constraints for both 2D and tilt-series data types in a single gradient descent optimization, and significantly improves the resolution of 3D reconstructions derived from such data.

Multi-particle refinement in *M* requires an initial tilt-series alignment, e. g. established using IMOD (*43*), as well as the particle poses, e. g. refined in RELION (*55*). At the beginning of each iteration, individual 2D particle tilt images are extracted from the original tilt-series. For extraction and subsequent refinement, 3D particle positions are related to their 2D positions in individual tilt images by applying the volume deformation model, the affine transform of the sample at the respective tilt angle, and the deformation model of the respective tilt image. The defocus value for each particle view is obtained by adding the Z coordinate of the transformed particle position to the global defocus value for the respective tilt image. Particle rotations are related to rotations in individual 2D images by applying the affine transform of the sample at the respective angle.

All per-particle parameters and the hyperparameters of the multi-particle system are optimized to maximize the cross-correlation between reference projections multiplied by the CTF and spectrally weighted according to accumulated dose and tilt angle, and the individual 2D particle tilt images summed over all particles and tilts. To achieve this in a gradient descent-like procedure, gradients must be calculated for all parameters with respect to the cost function in each optimization step. This calculation can be performed efficiently for particle poses using a finite differences numerical approach. However, most of the hyperparameters affect multiple particles, and a finite difference calculation must re-evaluate the cost function for all of these particles, making it computationally costly. Here, the same approach formulated in Warp (*48*) is employed: Before the optimization starts, partial derivatives are pre-computed for every affected 2D particle parameter with respect to each hyperparameter. During optimization, the hyperparameter gradients can then be efficiently calculated as a linear combination of the gradients of the affected 2D particle parameters.

The optimization is typically performed for 20 steps. At the end of the refinement of each tilt-series, individual 2D particle tilt images are back-projected into 3D volumes through Fourier-inversion (*56*), weighted by their local CTF and the dose- and angle-dependent spectral weights. Once refinement of all tilt-series is finished, reconstruction with Fourier-space gridding and amplitude correction is performed using the procedure described in RELION 1 (*55*) to obtain the final 3D maps. This approach, like emClarity (*53*), omits the lossy intermediate step of reconstructing individual sub-tomograms for each particle, and obtaining the final reconstructions by averaging these volumes, as implemented in other tools (*46*, *55*, *57*, *58*). The improved reconstructions can be used in another refinement iteration to further improve the resolution. As all parameters are optimized simultaneously in each iteration, the procedure's convergence is rather quick, taking 3 to 4 iterations. At last, sub-tomograms and their corresponding 3D CTF/wedge models can be re-generated after *M* refinement, with the updated hyperparameters taken into calculation inside Warp.

<u>Visualization and analysis of sub-tomogram analysis results</u>

RELION post-processing was used to calculate FSC and perform B factor sharpening. UCSF Chimera (*59*) and ChimeraX (*60*) were used for visualization and figure generation. Movies were generated using the "Volume Series" function in ChimeraX. Angular distribution maps were made by scripts described in cryo-EF (*61*). 3D FSC calculation and plots were performed in the online 3D FSC server (*50*). Sphericity values were calculated in 3D FSC analysis to measure directional resolution anisotropy resulting from preferential angular distribution. Quantification and plots were done in MATLAB (R2016b, The Mathworks, Inc. Massachusetts, USA).

<u>Homology modeling and structure prediction</u>

Sequence alignments were generated using clustal omega (*62*), disorder prediction was performed with IUPred2A (*63*) and secondary structure prediction using Jpred (*64*). Results were visualized in JalView (*65*).

The *M. pneumoniae* ribosome homology model was generated in SwissModell (*66*) using the *B. subtilis* MfiM-stalled ribosome (PDB 3J9W) (*67*) as a template for the ribosomal proteins, while retaining the rRNAs from *B. subtilis*. The validity of the homology models was verified by

CLMS, as well as using rigid-body fitting into the available densities (Fig. S9). Independent structural validation of crosslink data quality was performed by mapping crosslinks on homology models of additional complexes detected by CLMS, yielding a good agreement between crosslinks and predicted structures (Fig. S3), further confirming the native state of complexes in DSS- and DSSO-treated cells. For proteins L22 and L29, CLMS information was also used to guide the assignment of structured regions unique to *M. pneumoniae* that were observed in the density (Fig. S10-11). Disordered regions not localised in the density and presenting large crosslink violations were removed from the model. For Cm-treated cells, the ribosome homology model was based on the *T. thermophilus* chloramphenicol-bound ribosome (PDB 6ND5). The state of the ribosome in the stalled expressome from PUM-treated cells was determined by observation of the position of the L1 stalk, of EF-G and tRNA states (*68*). The homology model was based on the *E. coli* pre-translocation ribosome (*28*), PDB 4V7D).

Homology models of the *M. pneumoniae* RNAP were generated in SwissModel using the NusA-bound paused elongation complex from *E. coli* as template (PDB code 6FLQ, (*22*). Highly flexible loops and areas not covered in the sequence alignment were removed from the model and kept fully flexible in subsequent modeling steps (Table S4, "fully flexible coarse-grained regions"; Fig. S15). The NusG model was generated in SwissModel using NusG in the *E. coli* elongating RNAP complex as template (PDB 6C6U, (*69*), and placed relative to the RNAP accordingly, generating a combined RNAP-NusG model.

The homology models of the ribosome and RNAP were fitted into densities for the RNAP-ribosome complex, ribosome alone and the PUM-stalled complex using the fitmap tool in UCSF Chimera (*59*). A homology model of NusA was generated using MODELLER (*70*) based on an alignment with known NusA structures for the N-terminal, S1 and KH domains in multi-domain assembler (MDA) (*71*). Due to the low sequence homology and predicted disorder, residues 366-540 were left fully flexible and not modelled as domains (see Supplementary text, Fig. S18). Additionally, a homology model for the structured region (residues 1-80) of the fermicute-specific RNAP δ subunit, observed crosslinked to the RNAP core in a region with unassigned density, was generated using MODELLER. Homology model accuracy was assessed as reported in Table S4.

Integrative structure modeling: representation and sampling

The building blocks described above and in Table S4 were used to build an integrative model of the *M. pneumoniae* RNAP-ribosome supercomplex using the integrative modeling platform (IMP, version 2.12) (*24*). The overall modeling workflow is described in Fig. S19. Prior to modeling, RNAP-NusG and the ribosome were fitted into densities and kept fixed in subsequent steps. Unambiguity of fit was assessed by deriving a Benjamini-Hochberg-corrected p-value for the fit as in (*72*). The p-value for the ribosome fit to the expressome density is $7.2e^{-13}$, while the p-value for the RNAP density to the multibody-refined expressome density is $5.02e^{-13}$ (N=100,000).

In a first step, the models were coarse-grained as rigid bodies comprising the top of the 30S subunit (S2, S3, S4, S8, S9 and S10), the NusG-bound RNAP, each of the NusA domains and the RNAP δ subunit. Within each rigid body, regions that were previously removed (i.e. disordered and flexible regions, as well as regions for which no homology model could be generated) were coarse grained as beads and kept fully flexible, which allowed unassigned density in the RNAP and NusG regions to be filled according to cryo-EM and CLMS information. The rigid body boundaries and coarse-graining level are reported in Tables S4 and S5.

The restraints were then encoded as follows: 486 DSS and 145 DSSO-derived crosslinks were represented as distance restraints using the Bayesian CrossLinkingMassSpectrometryRestraint function in IMP.pmi, with psi (crosslink nuisance) to be sampled. The upper distance limits were set to 30 Å and 28.9 Å Cα-Cα distance for DSS and DSSO, respectively. The cryo-EM density from multi-body refinement comprising the RNAP and NusA regions of the complex was approximated as a gaussian mixture model (GMM), using 700 gaussians (cross-correlation 0.885) in IMP, and encoded as a Bayesian restraint using the IMP.BayesianEM function (*73*). The theoretical densities for the rigid bodies and flexible regions used in the modeling were also represented as GMMs, with numbers of residues per gaussians reported in Table S5. Upon analysis of the density, cryo-EM fitting was not imposed for one RNAP α-CTD, as well as the coarse grained residues 366-540 of NusA. In addition, excluded volume restraints, at a resolution of 10 residues, and connectivity restraints were added to the scoring function. The weight of CLMS, connectivity and excluded volume restraints was set to 2.

Sampling was performed by Replica Exchange Gibbs sampling in IMP 2.12, using 24 replicas in a temperature range between 1.0 and 80 in 20 independent runs with randomised initial configurations. A model was saved every 10 sampling steps, with each sampling step allowing for a 0.1 radians rotation and a 6 Å translation of each bead and rigid body. The number of frames was set to 15,000. During the calculations, the RNAP core, comprising the non-coarse-grained regions of RNAP α NTDs, β (excluding flat-tip helix), β' and NusG, as well as ribosomal proteins (Table S4) were kept fixed, as they could be fit by prior rigid body fitting into the density.

Integrative structure modeling: scoring and analysis

The scoring was performed as reported previously (*74*, *75*). A subset of 4,624 models of the sampled 7,200,000 configurations generated in replica exchange sampling was selected based on cutoffs in crosslink satisfaction (minimum 90% crosslinks < 35 Å distance), BayesianEM score and excluded volume score, as described in Table S6. In order to assess sampling completeness, the solutions were split into two random samples, which were then compared to each other (Fig. S20B). The sampling precision was determined by rmsd-based clustering of NusA in the models at different thresholds (Fig. S20C), leading to an overall sampling precision estimate of 10.8 Å. Here, we defined sampling precision as in (*75*), i.e. as the r.m.s.d. threshold at which models are no longer different from each other in a statistically significant manner ($p>0.05$), the effect size is small ($V<0.1$) and more than 80% of models fall in clusters in the subsequent rmsd-based clustering procedures. The solutions were clustered based on Cα root-mean-squared-deviation using a 10.8 Å cutoff, derived from the overall sampling precision. The precision of each cluster was defined as the average distance to the solution closest to the cluster center, i.e. to the centroid model. The top-populated cluster precision is 6.43 Å and the cluster comprises 96% of selected models (Fig. S20D). At the clustering threshold derived from sampling precision, only one cluster of solutions is found. The centroid model, with the associated localization probability densities, is taken as the representative model for the integrative modeling solutions. Localization probability densities were defined as the probability of any voxel (here, 5x5x5 $Å^3$) being occupied by a specific region in model densities over the selected cluster, each of which is obtained by convolving superposed models with a Gaussian kernel (here, with a standard deviation of 10.0 Å). Models were visualised using PyMOL, UCSF Chimera (*59*) and UCSF ChimeraX (*60*). Crosslinks were visualized with XlinkAnalyzer (*76*).

### Supplementary Text

Crosslinking-derived protein interaction network

The reduced genome of *M. pneumoniae* is of the simplest known self-replicating organisms and has therefore become an important model for bacterial systems biology (*13*, *14*). The incomplete knowledge of the *M. pneumoniae* protein interaction hampers its use as a model organism in synthetic and systems biology. A major advantage of in-cell crosslinking over previous affinity-purification mass spectrometry (AP-MS) screens (*13*)**,** is that it preserves the cell's intrinsic heterogeneity and complexity, such as low-affinity interactors, that can be lost when working with lysates. Here, 53% of the identified PPIs identified with in-cell CLMS have previous evidence in *M. pneumoniae* or homologes, as covered by the STRING database or reported by AP-MS in *M. pneumoniae (13)* (Fig. S1C). We identified interactions between the subunits of well-annotated complexes including the ribosome, ATP synthase, DNA gyrase and the condensin complex (see Table S1 and Fig. S2). For the ribosome, we found several interactions with known associated proteins, including initiation factors IF-1 and IF-3, elongation factor G, and trigger factor. Moreover, ribosomal protein S10 (NusE) was crosslinked to the transcription antitermination factor NusB, an interaction previously reported in *E. coli* (*77*) (Fig. S2A).

The detection of crosslinks is biased towards the abundant proteome due to the relative low abundance of crosslinked peptides compared to their unmodified counterparts (*78*). Accordingly, interactions with highly abundant proteins, such as the chaperone proteins DnaK, Tuf and GroEL, account for 150 of the binary PPIs identified (26%) and appear as "hubs" in the network. Other proteins that have more than 15 unique PPIs each are enolase, glyceraldehyde-3-phosphate dehydrogenase, the uncharacterized integral membrane protein Y376 (Mpn376), phosphotransacetylase, pyruvate dehydrogenase E1 component subunit alpha, the uncharacterized lipoprotein Y052 (Mpn052), and L-lactate dehydrogenase. The interactions in which these proteins participate account for another 155 PPIs (27%) identified and are all in the top order of magnitude of protein abundance (Table S1). Many of these proteins are commonly found in AP-MS studies, which is partially related to their high abundance, and are labelled as contaminants there (*79*). It is intriguing that we find these proteins to engage in many protein-protein interactions *in situ*. This rules out the possibility that those frequent AP-MS observations are an artifact of the experimental conditions, such as cell lysis. Whether they have any specific function, other than being a component of quinary interactions in a crowded cytoplasm, remains to be seen.

In-cell CLMS can identify PPIs involving membrane proteins and other difficult-to-solubilise parts of the proteome. This is an advantage over techniques that are biased towards soluble complexes, such as AP-MS, two-hybrid screens or coelution analyses that currently populate protein interaction databases. Lipidated proteins or proteins with transmembrane domains were involved in 41% of the newly identified interactions (interactions not annotated in the STRING database (*80*), see Fig. S1). These include interactions between the high molecular weight cytadherence proteins Hmw1-Hmw2 (29 crosslinked residue pairs) and Hmw2-Hmw3 (26 crosslinked residue pairs), for which there are no structural models. *M. pneumoniae* possesses reduced biosynthetic capacity, as these bacteria rely on the uptake of most building blocks for metabolism. Accordingly, we detected ABC transporter complexes, which transport peptides (OppA, OppC, OppF) and phosphate (PstA, PstB, PstS, PhoU) (Fig. S2B). RNA turnover is important for controlling gene expression in bacteria (*81*). Many bacteria contain the exonucleolytic 5'-3' RNase J. In *B. subtilis* and other bacteria, this RNase consists of a heterotetramer of two paralogous enzymes, RNase J1 and J2. For *M. pneumoniae*, so far only one RNase J protein had been annotated. This protein interacts with the membrane-bound protein MPN621 which we identify as the missing paralog of RNase J (Fig. S2B). Moreover, the essential membrane-bound endoribonuclease RNase Y crosslinks to the essential membrane protein MPN262 (8 crosslinked residue pairs), which is specific for *M. pneumoniae* and its close relative *M. genitalium*, suggesting they form a complex (Fig. S2B).

In total, the crosslinking-derived protein interaction network contains 301 proteins of which 76 are functionally only poorly annotated or not at all. Our newly identified interactions place these proteins in a functional context. The topological modeling of their interactions based on CLMS data may aid the design of further experiments to understand their function in greater depth.

Sub-tomogram analysis at sub-nanometer resolution of in-cell structures

Due to its small cell size, 70%-90% of a *M. pneumoniae* cell could be sampled in one tomogram acquired at a pixel size of 1.7 Å/pixel. Approximately 300 ribosomes (including both 50S and 70S) could be localized in one tomogram, which equals to 300 to 500 ribosomes per *M. pneumoniae* cell, higher than those reported previously (*13*, *82*). By comparing the proportion of ribosomes following classification with the correlation coefficient results from template matching (Fig. S5G), we found that the percentage of 70S ribosomes decreased with the correlation coefficient, and at the threshold used here for peak extraction from template matching (400 top scores), about 90% of all ribosomes could be retrieved. Therefore, the current sub-tomogram dataset retrieved the majority of 70S ribosomes inside the cell.

Maximum-likelihood classification and other functions implemented in RELION enabled resolving the large heterogeneity and flexibility of in-cell ribosomes and associated complexes. The obtained 5.6 Å 70S ribosome density following Warp/*M* refinement (described in detail in the materials and methods) shows the potential of this newly introduced sub-tomogram analysis workflow for resolving macromolecular structures at the level of secondary structures.

It is of note that expressomes can only form for the ‘leading’ ribosome, whereas most (all trailing) ribosomes that are part of polysomes on highly expressed mRNAs cannot directly interact with RNAP. We captured ~3,000 well-resolved elongating expressomes that may represent one of the relatively stable conformations during transcription-translation coupling, while there might be other more flexible states that could not be structurally characterized. The 17,000 ribosomes with additional density at the mRNA entry site, which could not be resolved further, might contain these more structurally flexible and dynamic expressomes. As shown by the multi-body refinement results, the relative movement of RNAP to the ribosome is large and continuous (Fig. S14 and Movie 1). Multi-body refinement of the 17,000 ribosomes with additional density did not produce better resolved structures, nor did further classification runs with different settings (local masks, class number, angular search range, etc.), suggesting that these sub-tomograms may have both high compositional heterogeneity and large conformational flexibility. Stalling RNAP by PUM reduced the heterogeneity and flexibility, therefore a substantially larger fraction of 70S ribosomes were classified and structurally resolved as RNAP-ribosome supercomplexes in the PUM-treated dataset (Fig. 4). Moreover, disruption or alteration of the supercomplex architecture following either Cm or PUM treatment indicated that assembly of the elongating expressome might be highly dynamic and transient inside the cell. This is further exemplified by the dramatic shifts observed in ribosome populations following relatively short perturbation time (~15 minutes).

Aspects of the model

NusA, a ubiquitous transcription elongation factor of bacteria, interacts with nascent mRNA upon binding to the elongating RNAP. In *E. coli*, NusA has an N-terminal domain, an S1 domain, two KH domains and two terminal acidic-repeat (AR) domains (Fig. S18). The S1-KH1-KH2 (SKK) domains interact with the nascent mRNA in several biological processes, ranging from transcriptional pausing (*22*), to antitermination (*83*) and intrinsic termination, and is involved in synchronising transcription and translation (*4*). In *E. coli*, the second AR domain has been found to bind to the RNAP α-CTDs (*22*, *84*), and to interact with NusG *in vitro* (*85*). The AR domains are not present in *B. subtilis* NusA, which terminates after KH2. Similarly, *M. pneumoniae* NusA has a low-complexity region in place of AR2, and low sequence and charge conservation for AR1, which indicates that the fold and function of this region may not be conserved (Fig. S18A). In this study, we chose to employ homology models for the *M. pneumoniae* NusA NTD and SKK domains, and to fully coarse-grain regions after the second KH domain (Fig. S18, S22 and Table

S4). Moreover, due to their predicted disorder, these regions are not fitted to the cryo-EM density, but rather left fully flexible.

In the cryo-EM map of the active expressome from untreated cells, a density corresponding to a small domain connected by a long stretch of density connecting back to the RNAP α subunit N-terminal domain was observed (Fig. S18E). We suggest that this density does not belong to the C-terminal region of NusA due to its predicted disordered nature and the lack of crosslinks between the C-terminal region of NusA and RNAP. Thus, one of the α-CTDs is also represented as gaussians, and is used to fit the cryo-EM density in the IMP protocol. In the cluster center solution selected by the protocol, this α-CTD is positioned above the second KH domain of NusA and occupies the density connecting NusA KH2 and the α subunit core (Fig. S22).

The role of the essential factor NusA in transcription elongation, termination and antitermination has been investigated in a number of studies (*1*, *22*, *29*). In the *E. coli* λ-induced antitermination complex, NusA works in conjunction with NusB, NusE (S10), the λN protein and mRNA to form a stable complex that presumably displaces NusA from the nascent mRNA, preventing the formation of intrinsic termination hairpins and reprogramming the role of the NusA NTD from termination-promoting to antitermination (*83*, *86*). Antitermination in the *E. coli rrn* operon, encoding for rRNA genes, is also thought to occur via a mechanism in which NusB-NusE modulate NusA activity (*77*) in conjunction with S4 (*29*). Here, we do not find evidence for S10 or NusB being directly bound to NusA in the RNAP-NusG-NusA-ribosome supercomplex by cryo-EM. Indeed, despite a crosslink being observed between S10 and the NusA C-terminal region, there is no density available to accommodate S10 or NusB in contact with NusA. Moreover, the position of NusG in the elongating expressome precludes an interaction with S10 (Fig. S17C). This interaction may however still occur in the initial ribosome recruitment to RNAP, or in other transcription-translation coupling modes, as the residues involved are conserved (Fig. S17A).

While NusA interacts with nascent mRNA via its SKK domains, the interaction between NusA and the RNAP has been proposed to occur via multiple mechanisms. The structure of the *his*-paused elongation complex suggests binding between one α-CTD and AR2, consistent with previous studies *(84, 87)*, and an interaction between the NTD and the second α-CTD, besides a direct interaction between the NTD and the β flat-tip helix also present in the antitermination complex (*22*, *86*). The resolution in this study is not sufficient to unambiguously position the flat-tip helix. However, its position, as indicated by CLMS, is consistent with its proposed interaction with NusA. Similarly, density for one α-CTD is observed in proximity of KH2, while the second α-CTD could not be localized to a high precision. However, no density is observed for a binding mode to the NusA NTD corresponding to the *E. coli* paused elongation structure. Nevertheless, the overall position of the NusA NTD relative to RNAP closely resembles that of the paused elongation complex (Fig. S21).

The model is also consistent with interactions between the mRNA and the SKK domains, which may guide the mRNA into the entry site on the 30S ribosome (Fig. S22E). It is unclear whether the novel interactions presented here between NusA KH domains and S2, S3, S4, S5 subunits (Fig. S22E) play a role in regulating transcription or translation rates, or whether these contacts are a consequence of NusA positioning on the nascent mRNA. Nevertheless, the highly flexible nature of the elongating expressome (Movie 1) indicates that the domain arrangement in this complex is highly dynamic, possibly influencing and being influenced by the nature of the DNA sequence being transcribed, as well as the rate at which transcription and translation are proceeding.

### Supplementary figures

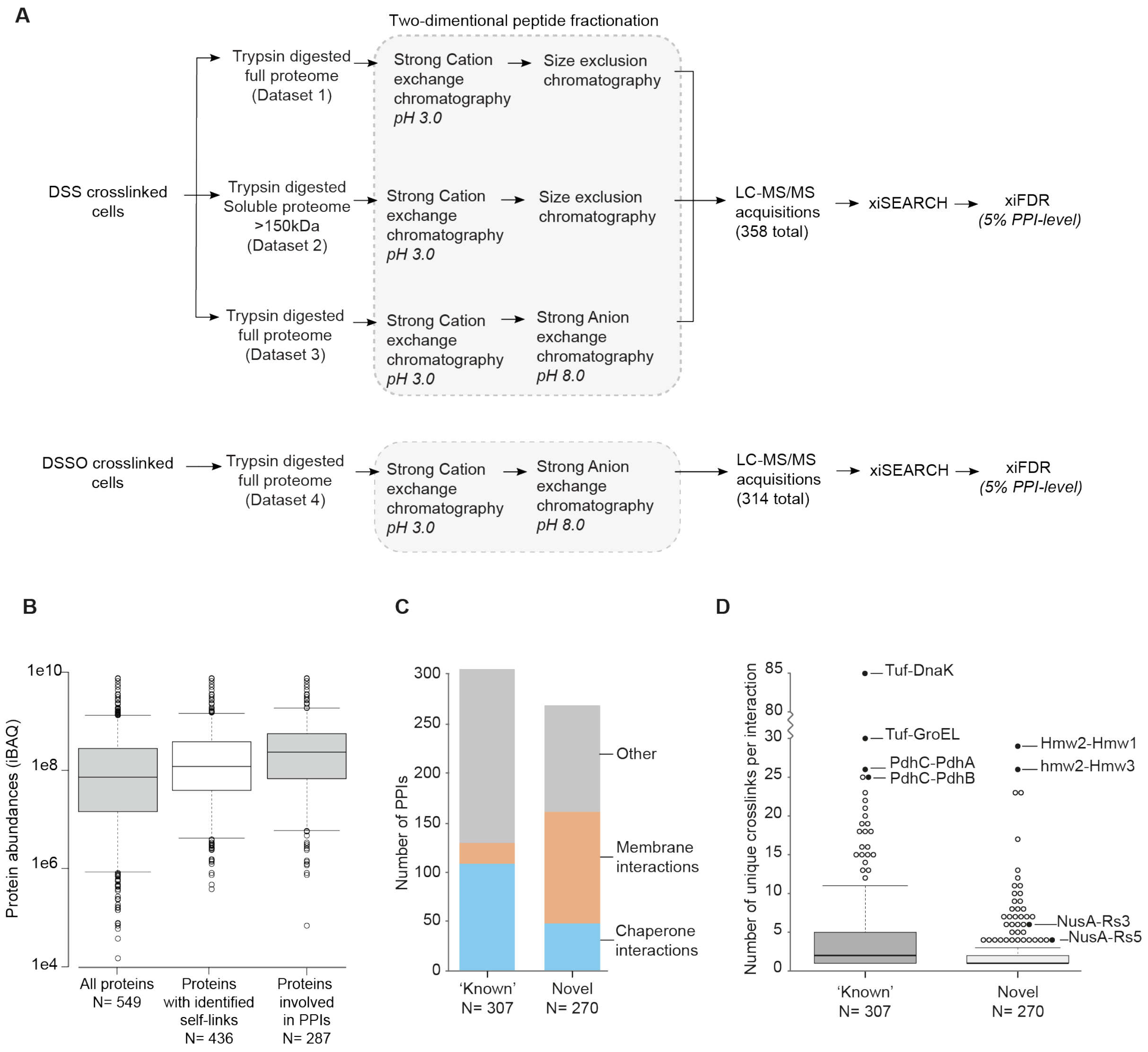


**Fig. S1.**

**Whole cell CLMS dataset.** (**A**) Cells were crosslinked with either DSS or DSSO and the crosslinked proteome was extracted and digested using Trypsin. For one dataset of DSS crosslinked cells, only the high molecular weight part of the soluble proteome (>150 kDa) was used (Dataset 2). Crosslinked peptides were sequentially enriched and fractionated in two dimensions by either strong cation exchange chromatography followed by size exclusion chromatography, or strong cation exchange chromatography followed by strong anion exchange chromatography. (**B**) 549 proteins were detected from *M. pneumoniae* and 83% of these were identified with CLMS. As crosslinked peptides are generally low-abundant in comparison to linear peptides, they tend to be detected in proteins present at higher copy numbers. The median intensity of proteins identified by shotgun proteomics is $7.4e^{7}$, but for proteins with identified intra-protein or inter-protein crosslinks it is $1.2e^{8}$ and $2.4e^{8}$ respectively. (**C**) More 'known' interactions than unknown interactions (according to STRING-DB v11 (*80*)) are reported. Of the 'novel' interactions, 59% involve either membrane associated proteins or chaperones. (**D**) Known interactions tend to have higher numbers of unique crosslinked residue pairs, suggesting higher abundance of these interactions/protein assemblies in the cell. Interactions between the high molecular weight cytadherence proteins (hmw2-hmw1; hmw2-hmw3) or NusA with 30S ribosomal (RS3 and RS5) proteins are not reported in STRING.

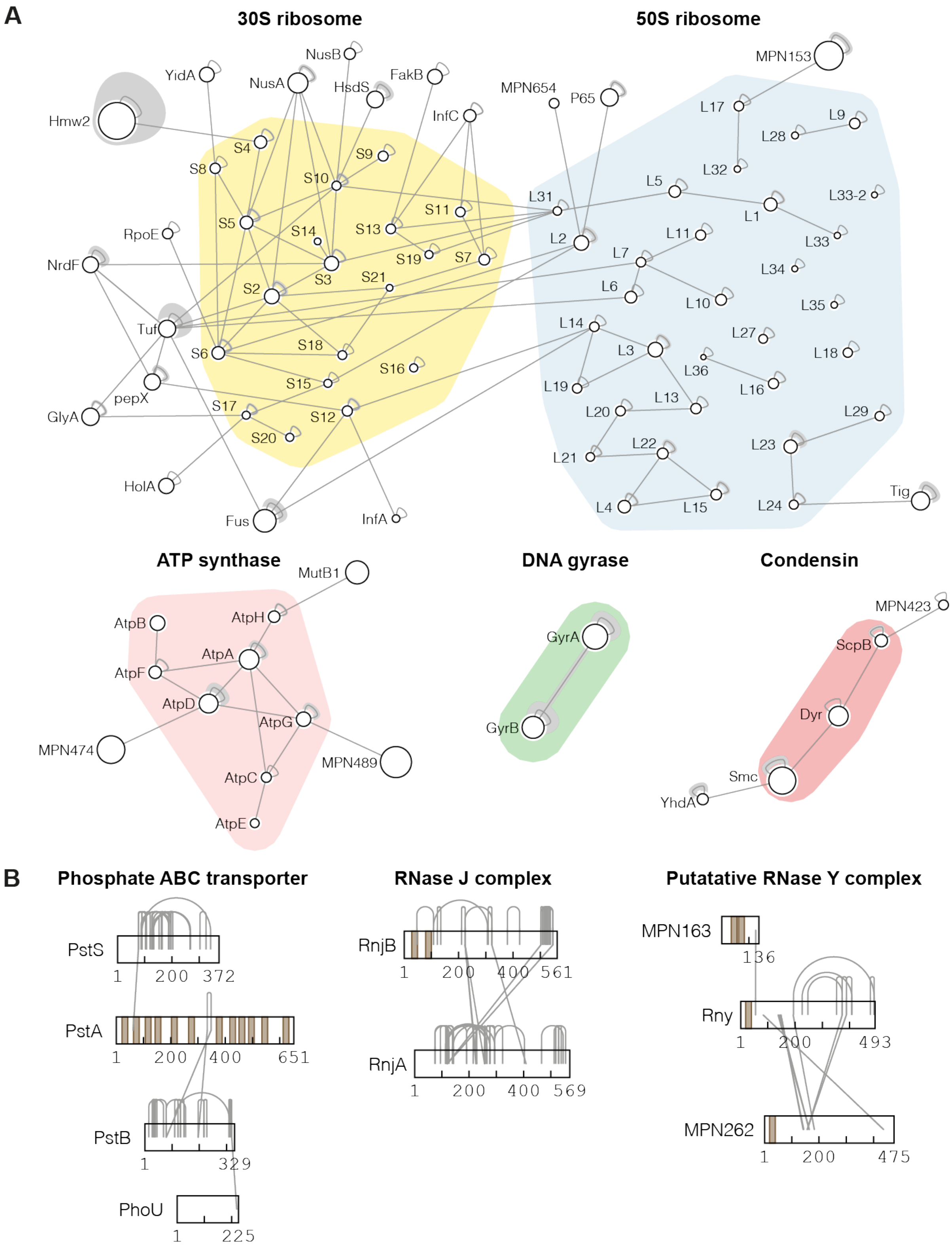


**Fig. S2.**

**Protein complexes and their interactors detected in the dataset.** (**A**) Networks showing all intra- and inter-protein crosslinks identified on known complexes. Any proteins found crosslinked to these are also shown. Circle diameter indicates relative protein size. Each edge represents one or more crosslinks. (**B**) Crosslink-based protein networks of previously uncharacterised interactions involving membrane-bound subunits. Brown lines on the sequence indicate transmembrane regions. The endoribonuclease RNase Y (Rny) does not have a previously annotated interaction partner.

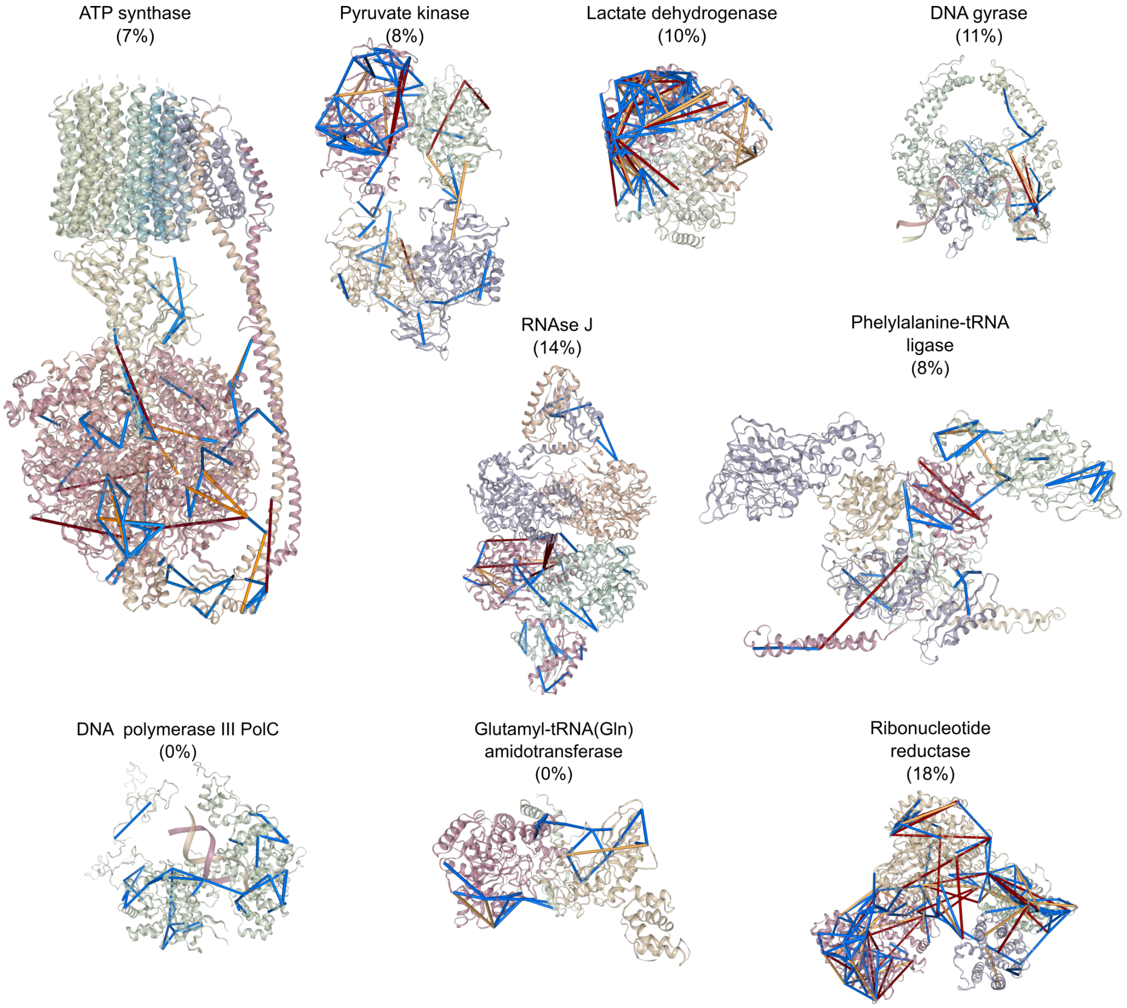


**Fig. S3.**

**Mapping of crosslinks on homology models of known complexes in *M. pneumoniae*.** Crosslinks mapped on homology models of detected protein complexes generated with SwissModel. Satisfied links (<28 Å) are displayed in blue, links near the violation threshold (28-35 Å) in yellow and violated links (>35 Å) in red. The percentage of violated links is shown in parentheses.

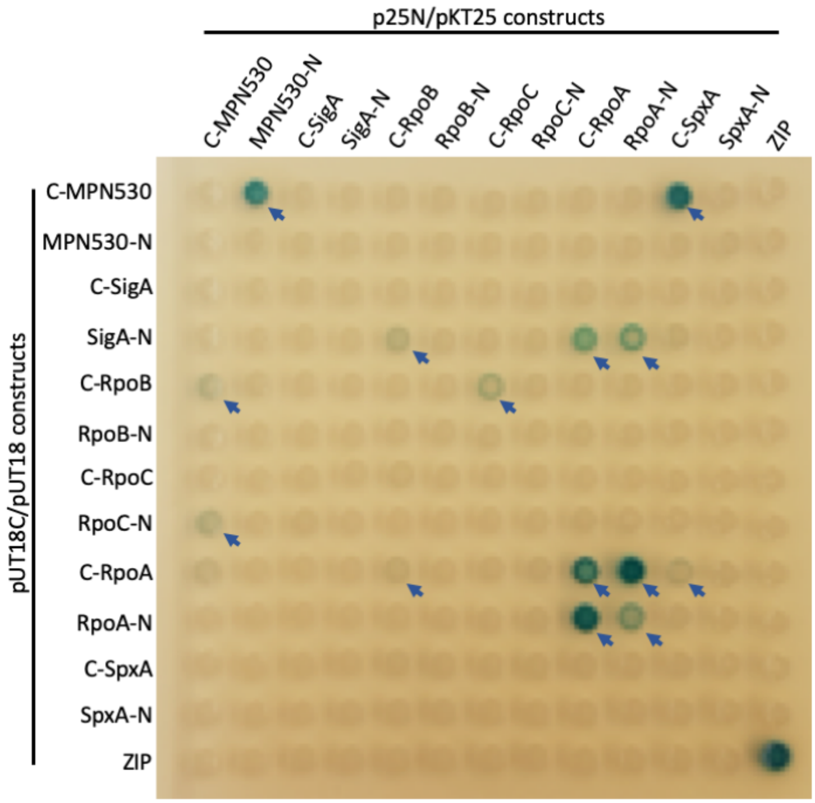


**Fig. S4.**

**Bacterial two hybrid screen confirms interaction of MPN530 and RNAP.** To confirm novel interactions identified by in-cell CLMS between MPN530 and the subunits β and β' of the RNAP, we studied the binary interactions by using a bacterial two-hybrid screen. In this system, interacting proteins reconstitute the *B. pertussis* adenylate cyclase, resulting in cAMP synthesis and subsequent activation of β-galactosidase synthesis. Colonies in which interactions are observed (β-galactosidase is expressed) are marked by blue arrows. MPN530 and SpxA showed a strong interaction. Moreover, MPN530 and RpoA exhibited strong self-interaction. We detected a weak, but significant, interaction between MPN530 and β (RpoB) as well as β' (RpoC) (left column). We could also confirm the interaction between SigA with α (RpoA) and β (RpoB) and the interaction of SpxA with α (*16*). Furthermore, as expected, all tested subunits of the RNA polymerase interact.

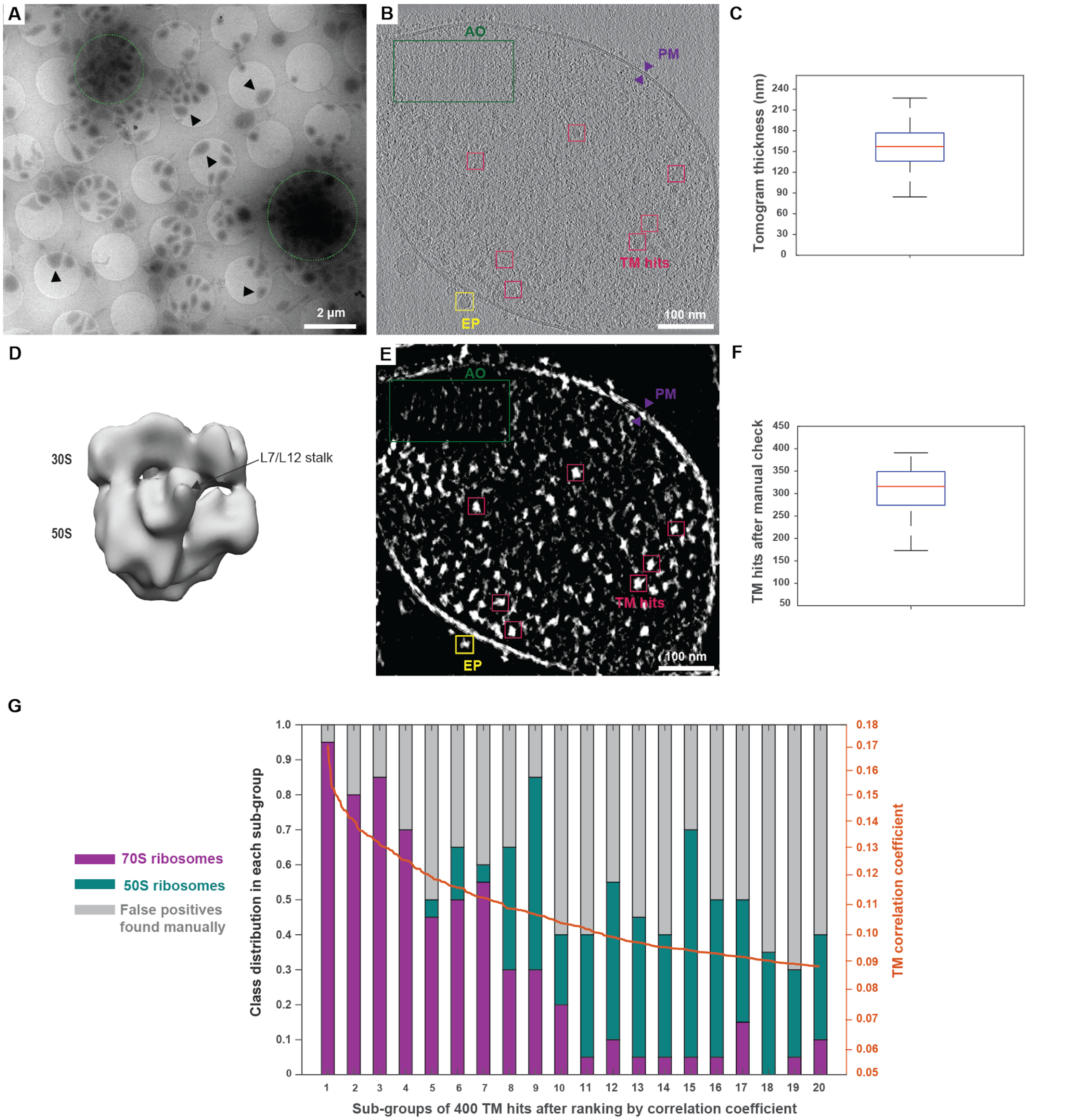


**Fig. S5.**

**Cryo-EM of *M. pneumoniae* and ribosome localization.** (**A**) Individual *M. pneumoniae* M129 cells (arrowheads) and cell colonies (green dashed circles) were grown on cryo-EM grids. cryo-ET data were collected on individual cells. (**B**) XY slice of a conventional defocus tomogram. Annotation of representative ribosomes localized by template matching (red boxes), the attachment organelle (AO, green box), the plasma membrane (PM, purple arrowheads) and unknown extracellular particles (EP, yellow boxes). (**C**) Distribution of cell thickness estimation from tomographic reconstructions. The thickness ranges from 84 nm to 284 nm (black bars), with a median value of 158 nm (red line). Blue box represents the 0.25-0.75 quartile. (**D**) *De novo* ribosome sub-tomogram average reconstructed from 404 manually picked ribosomes from VPP tomograms. Density was low-pass filtered to 30 Å and used as reference for template matching. (**E**) Cross correlation coefficients of the ribosome reference with the tomographic volume in (B). Brighter pixels indicate higher cross correlation values. Obvious false positives such as membranes, carbon edges and extracellular particles were manually excluded. (**F**) Number of ribosomes picked per tomogram after template matching and visual validation. The median number is 316 (red line), while most tomograms contained between 274 to 349 ribosomes (blue box). (**G**) The top 400 template matching hits from a representative tomogram were ranked by correlation coefficients (yellow curve) and divided into 20 sub-groups. After visual inspection and 3D classification of the sub-tomograms, percentages of obvious false positives (grey), 50S (green) and 70S (magenta) ribosomes within each sub-group were calculated and plotted, showing that a cutoff of 400 peaks retains most ribosomes in the data.

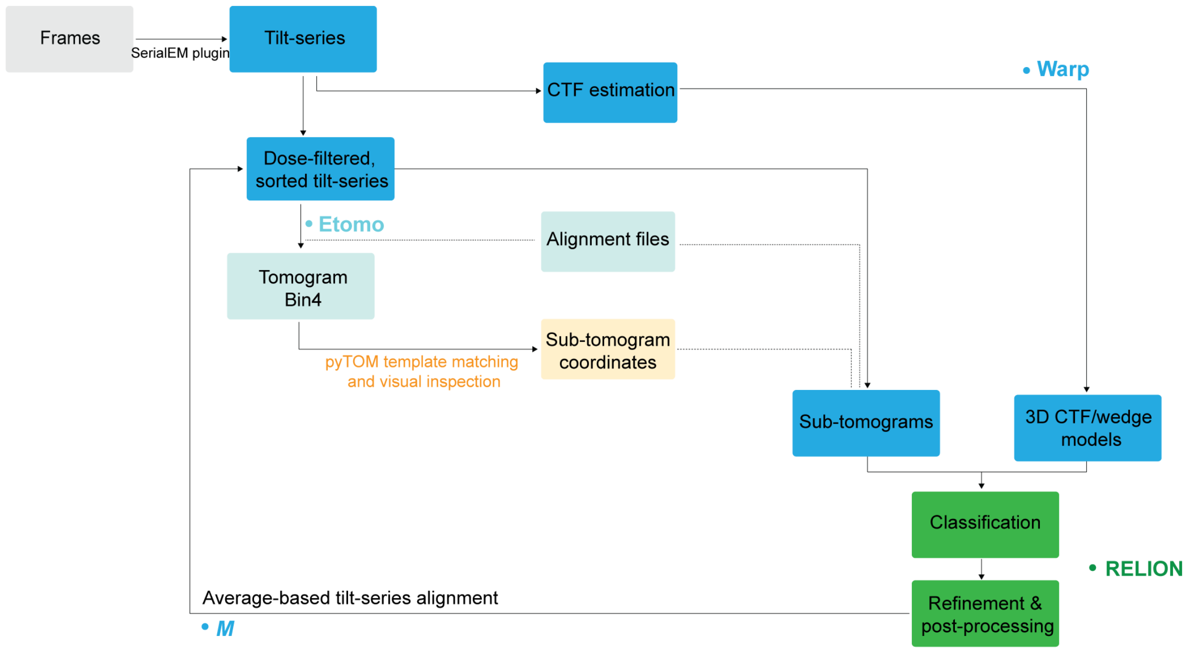


**Fig. S6.**

**Sub-tomogram analysis workflow.** Frames were aligned on-the-fly with the IMOD alignframes plugin in SerialEM. Dose-symmetric tilt-series were pre-processed in Warp for CTF estimation, sorting of the tilt images and dose filtering. Tilt-series were aligned in Etomo with 4x binning using gold fiducials and tomograms generated using back-projection. Template matching was performed in pyTOM, followed by visual inspection using MATLAB/TOM toolbox. Tilt-series, alignment files, CTF files and particle coordinates were utilized by Warp to reconstruct sub-tomograms, as well as per sub-tomogram 3D CTF/wedge models containing CTF, dose filtering and missing wedge information. Sub-tomograms were analysed in RELION 3.0, including 3D classification, 3D refinement, multi-body refinement and post-processing. RELION refinement and Warp processing results were used in *M* for 3D average-based (i.e. ribosomes) alignment for each tilt-series. Sub-tomograms and 3D CTF/wedge models can be re-generated after *M*, with the refined alignment parameters.

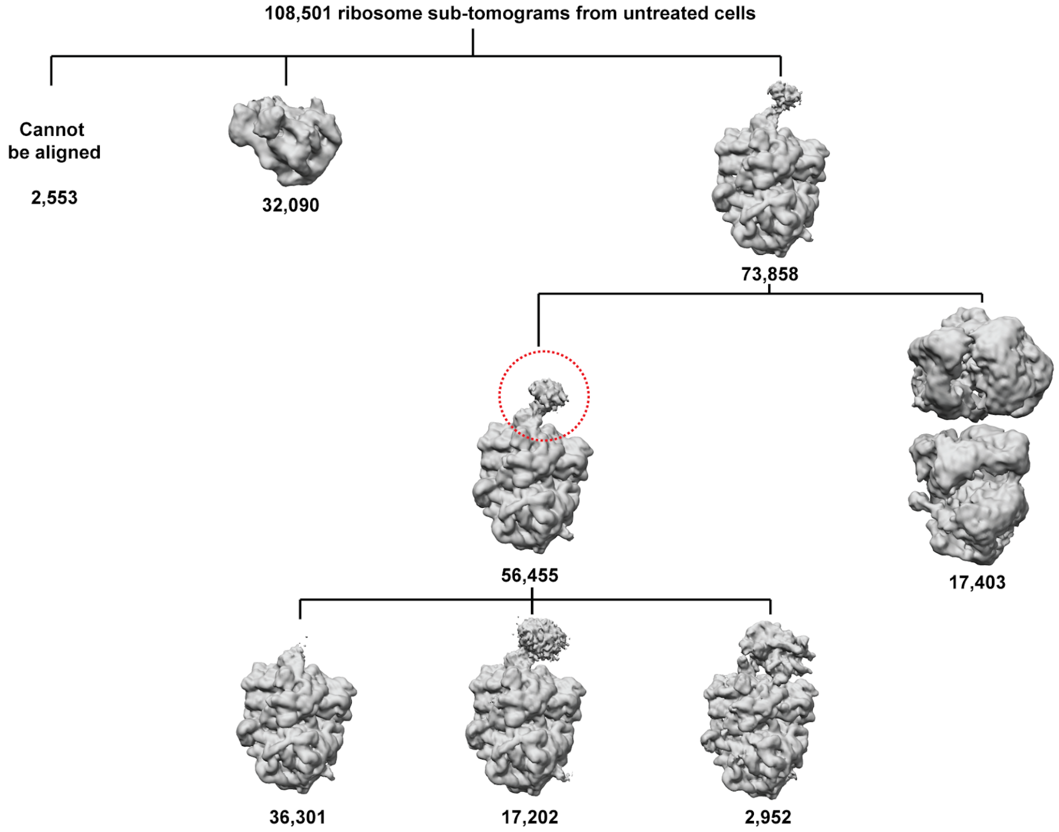


**Fig. S7.**

**Maximum likelihood-based classification of ribosome sub-tomograms from untreated *M. pneumoniae* cells.** In total, 108,501 ribosome sub-tomograms were reconstructed and then classified in RELION. Except for 2,553 sub-tomograms that could not be aligned, 2 distinct classes of ribosomes were identified: 50S ribosome (32,090) and 70S ribosomes (73,858). Prior to subsequent focused classification near the mRNA entry site (red dashed circle), 70S ribosomes in closely assembled polysomes (17,403) were sorted out. The rationale to exclude closely-assembled ribosomes is that neighboring ribosomes could interfere with further focused classification and are unlikely to interact with RNAP. 2,952 sub-tomograms exhibit a well-defined RNAP density near the mRNA entry site, while 36,301 ribosomes do not have significant additional density. For the class of 17,202 ribosomes, the additional density near mRNA entry site is significant and of similar size to a RNAP, but could not be further classified or refined. They may represent other, more flexible, supercomplexes formed during transcription-translation coupling.

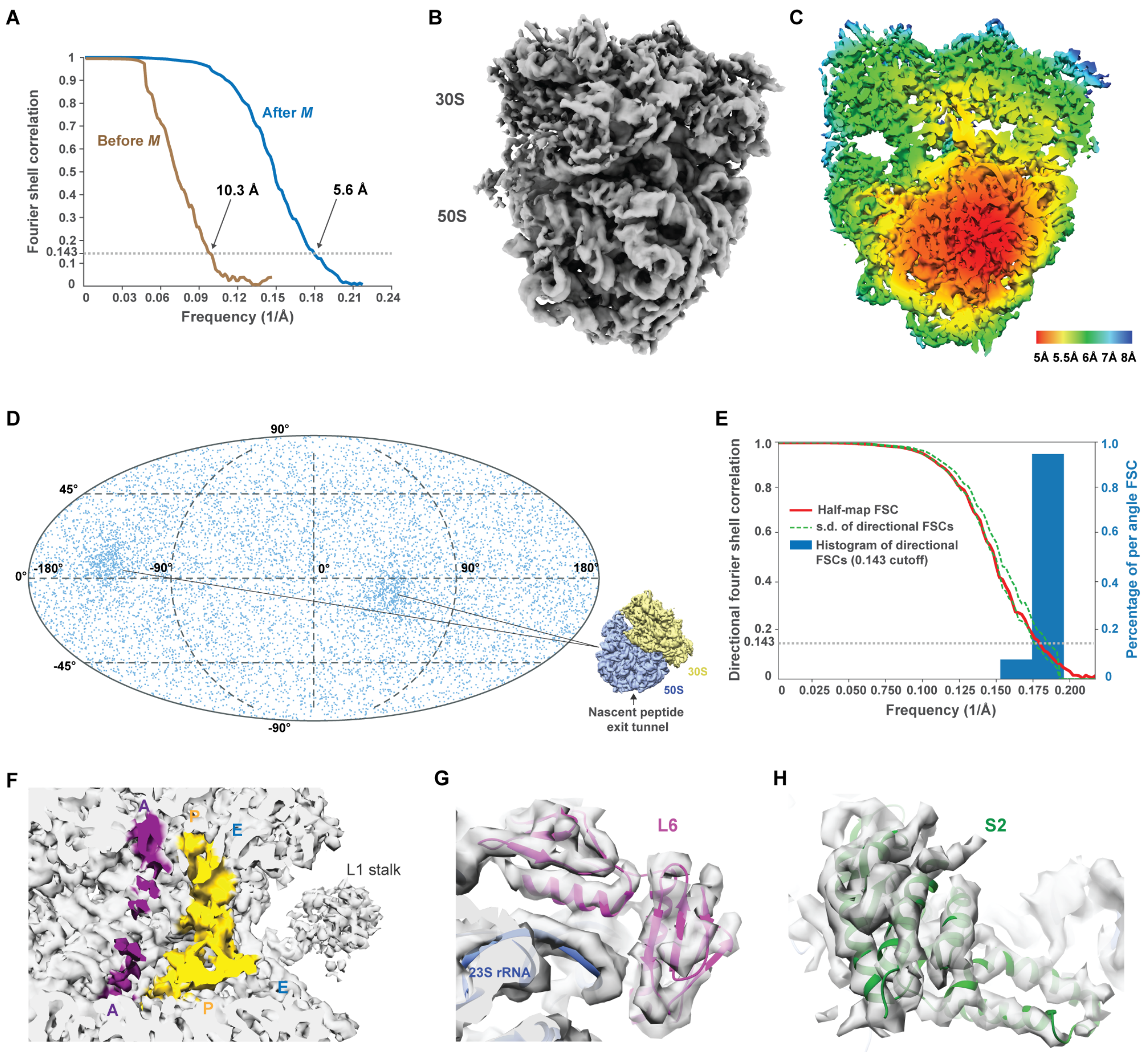


**Fig. S8.**

**5.6 Å in-cell *M. pneumoniae* ribosome structure obtained by sub-tomogram averaging.** (**A**) Refinement of all 70S ribosome sub-tomograms generated a map of 5.6 Å resolution (FSC=0.143). Refinement of the tilt-series in *M* resulted in a significant resolution improvement. (**B**) Surface representation of the 5.6 Å in-cell ribosome density. (**C**) Central slice of the local resolution map. (**D**) Angular distribution plot shows that the dataset covered all particle orientations, but exhibited weak preferential angular distribution. As indicated by the ribosome structure (bottom right), these two peaks represent ribosomes oriented parallel to nascent peptide exit tunnel in the data. (**E**) 3D FSC analysis. Calculated sphericity of 0.985 indicates minimal directional resolution anisotropy of the density. (**F**) Densities corresponding to tRNAs in A and P site. No significant density for E site was detected. (**G**, **H**) Homolog structures (based on PDB 3J9W) of two ribosomal proteins, L6 (**G**) and S2 (**H**), docked into the 70S density.

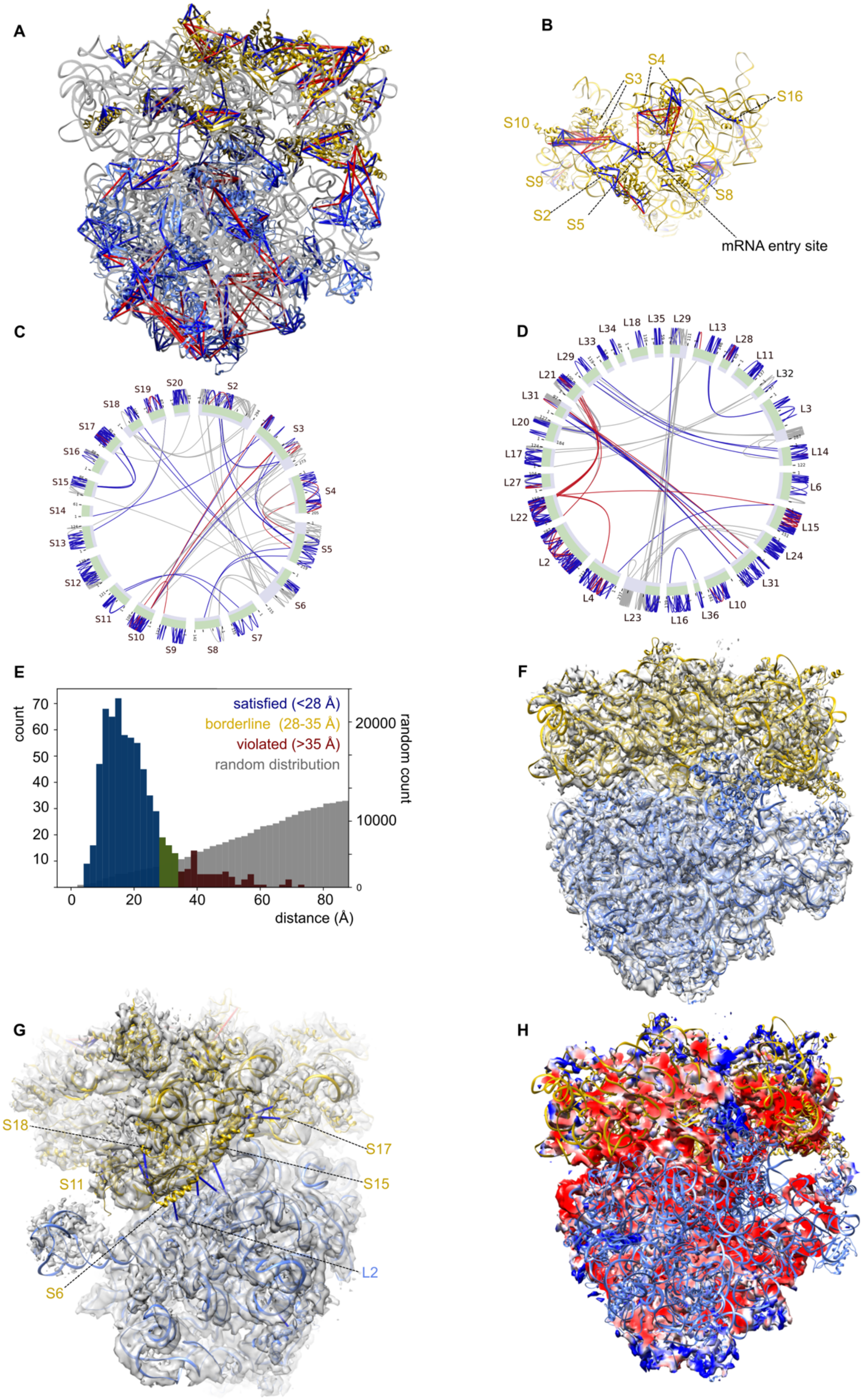
A
B
S4
S3
S16
S10
S9
S8
S2
S5
mRNA entry site
C
S19
S20
S2
S18
S17
S3
S16
S15
S4
S14
S5
S13
S12
S6
S11
S10
S9
S8
S7
D
L34
L18
L35
L29
L13
L33
L28
L29
L11
L21
L32
L31
L3
L20
L14
L17
L27
L6
L22
L15
L24
L2
L31
L4
L10
L23
L16
L36
E
count
satisfied (<28 Å)
borderline (28-35 Å)
violated (>35 Å)
random distribution
random count
distance (Å)
F
G
S18
S17
S11
S15
L2
S6
H

**Fig. S9.**

**The *M. pneumoniae* ribosome homology model**. (**A**) Crosslinks mapped on homology models of the ribosome based on the *B. subtilis* Mfim-stalled ribosome (PDB 3J9W) generated using Swissmodel. Subunits not found in the density (L1, L7/L12, L9 ,S21) and fully flexible regions have been removed. The rRNA from *B. subtilis* has been retained, as the resolution was not sufficient to build the *M. pneumoniae* rRNA into the map. The largest violations are observed in the peptide exit tunnel region (proteins L21, L22). Satisfied links (<35 Å) are displayed in blue, and violated links (>35 Å) in red. (**B**) View of the top of the 30S subunit in the homology model with crosslinks displayed labelled by distance as in (A). (**C**) Circle representation of the crosslinks on the 30S subunit labelled by distance as in (A). (**D**) Circle representation of the crosslinks on the 50S subunit labelled by distance as in (A). (**E**) Distribution of 683 crosslink distances obtained from the ribosome model compared to a theoretical random distribution of lysine-lysine and lysine-N-termini crosslinks from the same model. (**F**) Fitting of the *M. pneumoniae* ribosome homology model to the 5.6 Å in-cell ribosome density. (**G**) View of the 30S-50S interface with inter-subunit crosslinks and density displayed. (**H**) Cross-section of the in-cell ribosome density colored by local cross-correlation (CC) to a theoretical 6 Å density derived from the ribosome homology model. Blue: CC<0.7; White: 0.7<CC<0.9; Red: CC>0.9.

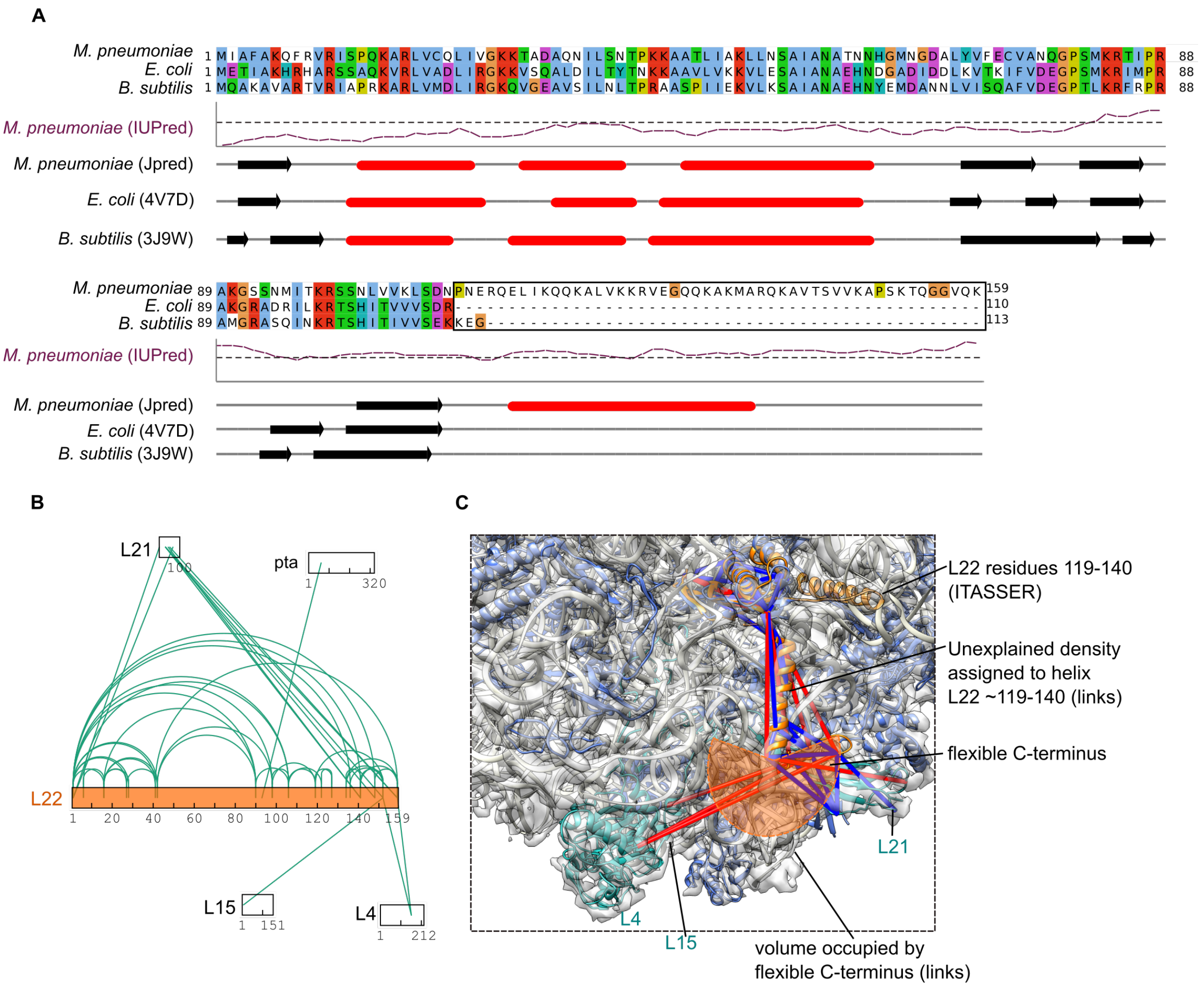


**Fig. S10.**

**The C-terminal region of ribosomal protein L22 in *M. pneumoniae*.** (**A**) Sequence alignment of *M. pneumoniae*, *E. coli*, and *B. subtilis* L22 proteins. For *E. coli and B. subtilis*, the secondary structure as shown in PDBs 4V7D and 3J9W is displayed. For *M. pneumoniae*, the secondary structure was predicted with Jpred. β-sheets are shown as arrows, α-helices as cylinders. The long-range disorder prediction of IUPred is also shown, with the threshold of 50% disorder likelihood displayed as a dashed line. *M. pneumoniae* presents a unique C-terminal extension from residue 113, not found in other structures. (**B**) The *in-cell* crosslinking network of L22. (**C**) Placement of the L22 C-terminal region (orange) according to in cell cryo-EM density and crosslink data. The original structure prediction for the L22 C-terminal region, obtained with I-TASSER (shown in transparent orange), is found outside the EM density map and did not agree with crosslinking information. Using crosslinks, we unambiguously assigned the unexplained density to the C-terminal region of L22, although the resolution was not high enough to determine helical orientation and boundaries. The density and a crosslink to L21 are consistent with a long α-helix up to residue ~140, located in proximity of the final hairpin of 23S rRNA domain I. Crosslinks between L22 and L21, L4, as well as L15 in residues ~141 to 159 are consistent with this region being highly flexible. Crosslinks shorter than 35 Å are shown in blue, while those longer than 35 Å are in red.

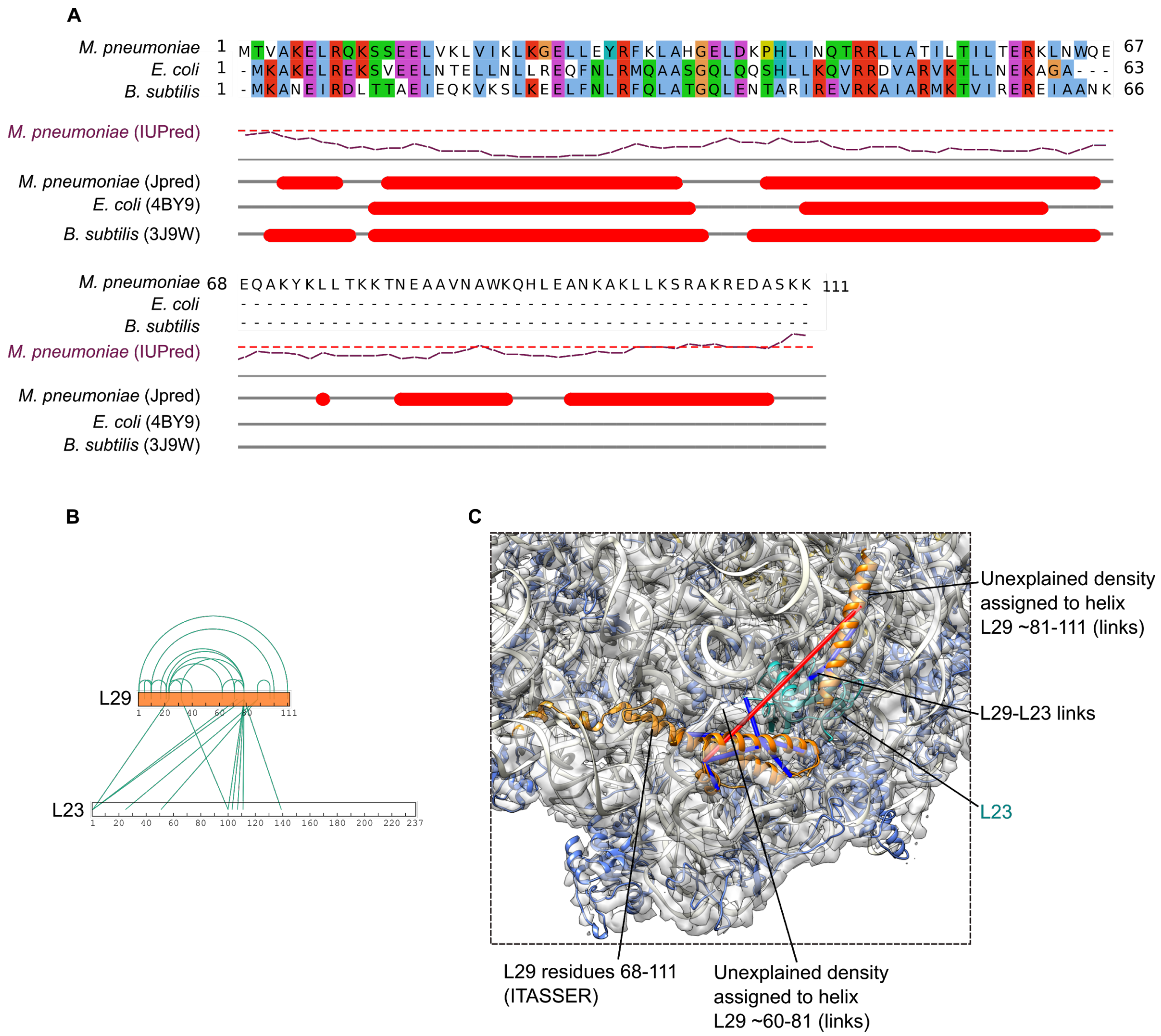


**Fig. S11.**
**The C-terminal region of ribosomal protein L29 in *M. pneumoniae*.** (**A**) Sequence alignment of *M. pneumoniae*, *E. coli*, and *B. subtilis* L29 proteins. For *E. coli and B. subtilis*, the secondary structure as observed in PDBs 4V7D and 3J9W is displayed. For *M. pneumoniae*, the secondary structure was predicted with Jpred. β-sheets are shown as arrows, α-helices as cylinders. The long-range disorder prediction of IUPred is also shown, with the threshold of 50% disorder likelihood displayed as a dashed line. *M. pneumoniae* presents a unique C-terminal extension from residue 68, not found in other structures. (**B**) The *in-cell* crosslinking network of L29, confirming its placement near L23. (**C**) Placement of the L29 (orange) C-terminal region according to the cryo-EM density and crosslink data. The original prediction for the L29 C-terminal region, obtained with I-TASSER (shown in transparent orange), lay outside the EM density map and did not agree with crosslinking information. Using crosslinks, we could assign the unexplained density to the C-terminal region of L29, although the resolution was not high enough to determine helical

orientation and boundaries. The density, secondary structure prediction and crosslinks to L23 are consistent with a loop connecting the N-terminal region of L29 to a long C-terminal α-helix up to residue ~111. Crosslinks shorter than 35 Å are shown in blue, while those longer than 35 Å are in red.

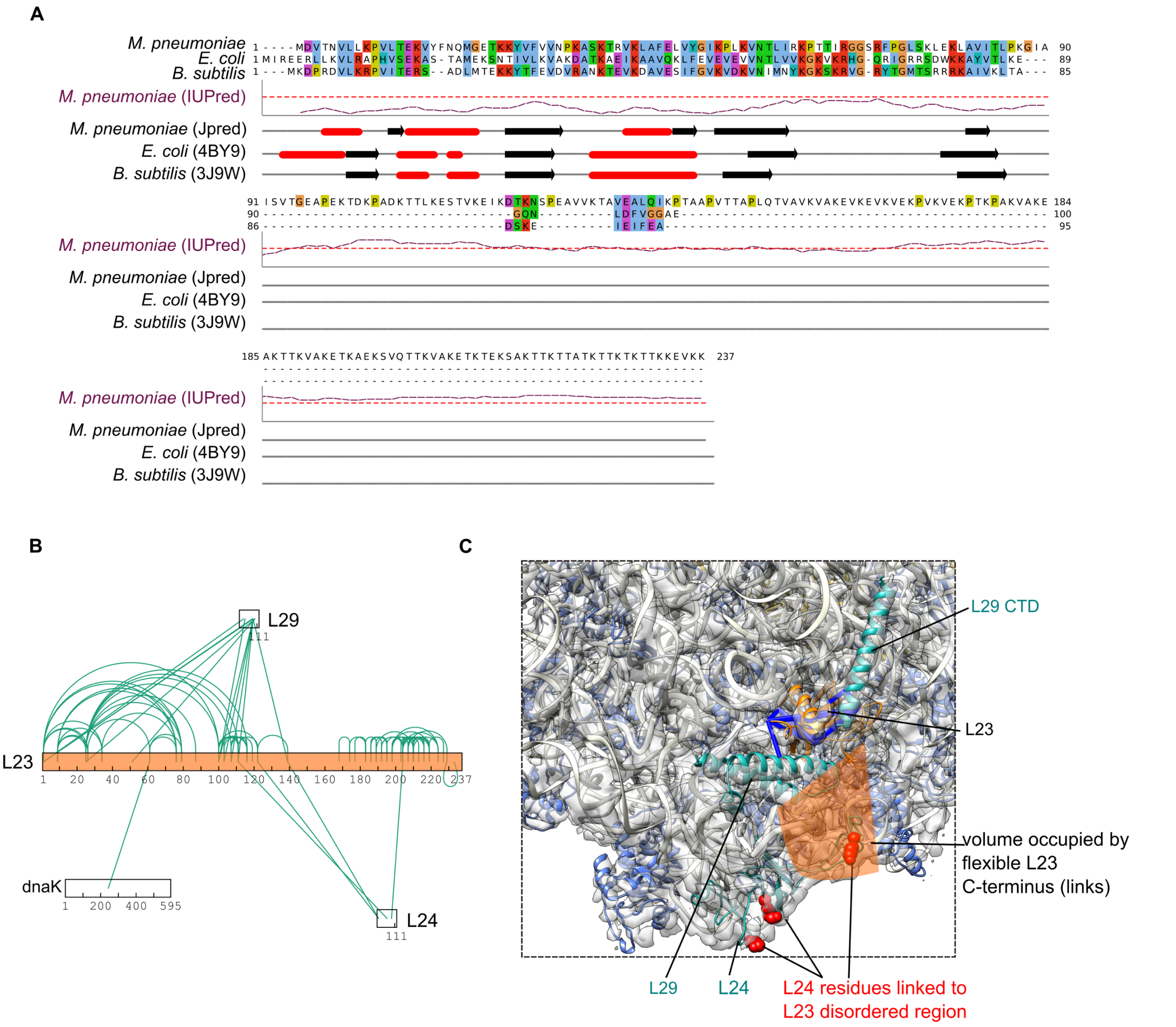


**Fig. S12.**

**The C-terminal region of ribosomal protein L23 in *M. pneumoniae*.** (**A**) Sequence alignment of *M. pneumoniae*, *E. coli*, and *B. subtilis* L23 proteins. For *E. coli and B. subtilis*, the secondary structure as observed in PDBs 4V7D and 3J9W is displayed. For *M. pneumoniae*, the secondary structure was predicted with Jpred. β-sheets are shown as arrows, α-helices as cylinders. The long-range disorder prediction of IUPred is also shown, with the threshold of 50% disorder likelihood displayed as a dashed line. L23 is predicted to be largely disordered from residue 94. (**B**) The *in-cell* crosslinking network of L23, which includes a crosslink to the highly abundant dnaK. (**C**) Placement of the L23 structured domain according to the cryo-EM density and crosslink data. The density, secondary structure prediction and crosslinks to L23 are consistent with a disordered C-terminus in the proximity of L24 and a structured N-terminal domain placed as in the *B. subtilis* ribosome. Accordingly, the L23 structured region lies near the L29 N-terminal domain and the L29 C-terminal helix found in *M. pneumoniae* (see Fig. S11). Crosslinks shorter than 35 Å are shown in blue, while those longer than 35 Å are in red. L24 residues cross-linked to the disordered region of L23 are shown as red spheres.

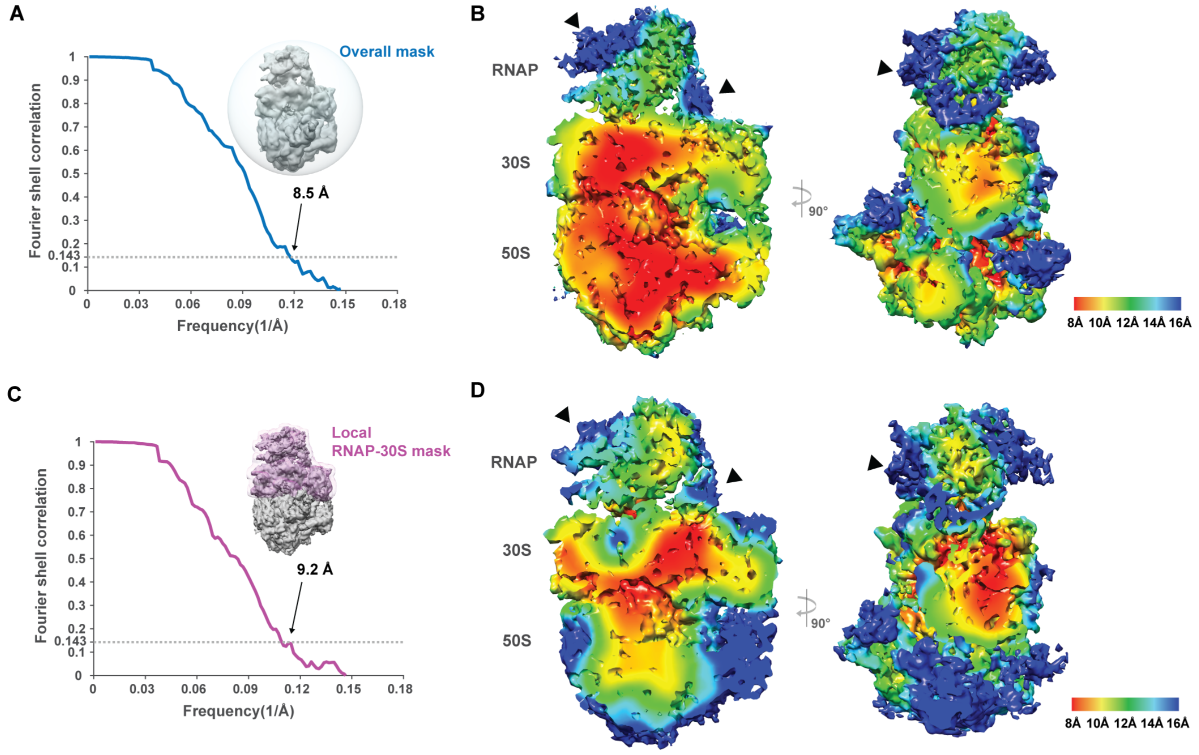


**Fig. S13.**

**In-cell RNAP-ribosome supercomplex determined at sub-nanometer resolution.** 2,952 sub-tomograms from the class representing RNAP-ribosome supercomplexes were refined with different masks. (**A**) Refinement with a spherical mask (insert). Global resolution of the refined density is 8.5 Å. (**B**) Local resolution map shows that the ribosome core has higher resolution, while the RNAP density is of relatively lower resolution, especially for the flexible clamp region and the bridging part between RNAP and the 30S (arrowheads). (**C**) Focused refinement with a RNAP-30S mask (insert) generated a map of overall resolution 9.2 Å. (**D**) 30S is of higher local resolution, while the 50S is blurred, likely due to ratcheting. Local resolution for RNAP improved compared to the refinement with a spherical mask (B).

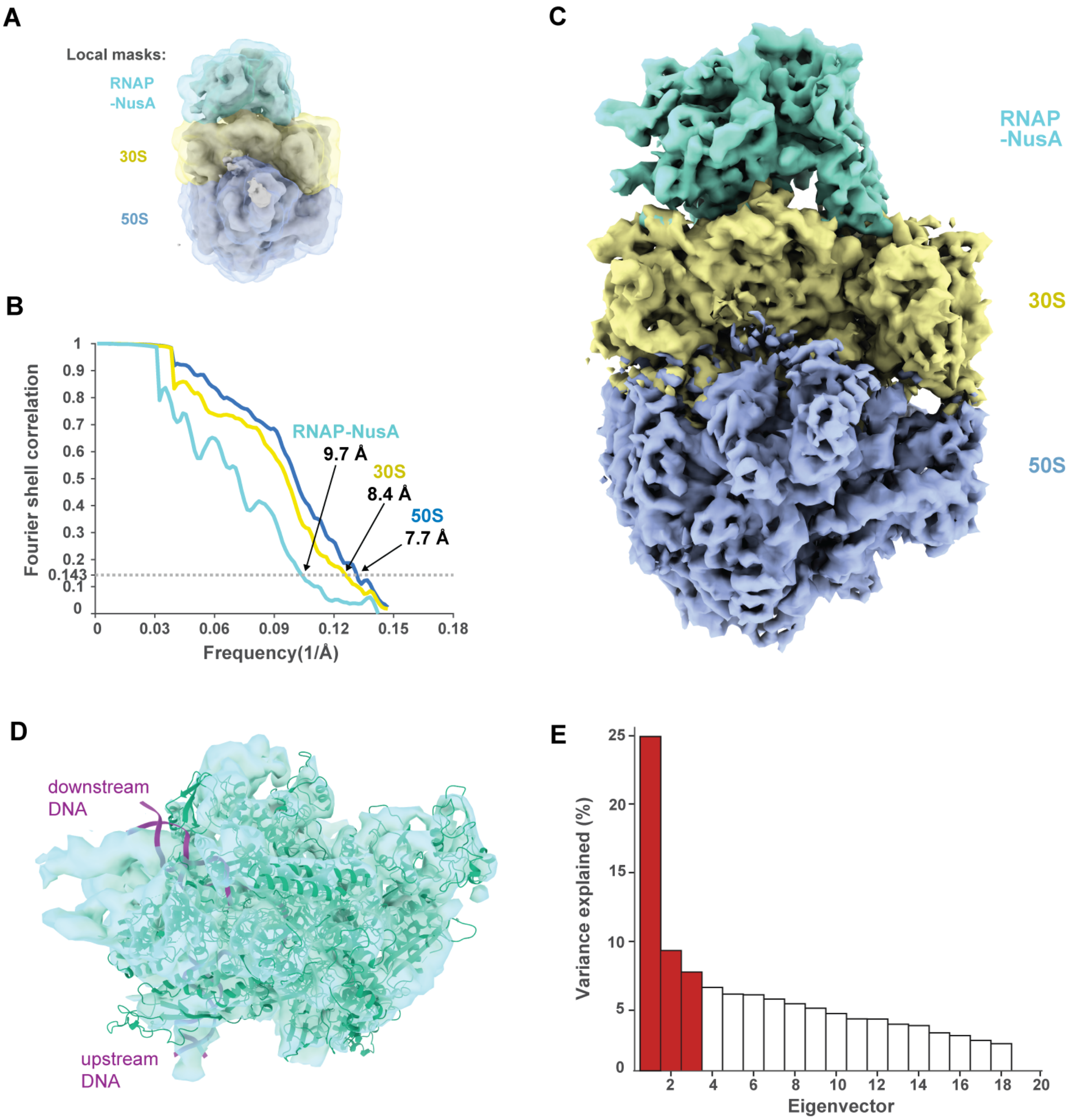


**Fig. S14.**

**Multi-body refinement reveals high conformational flexibility of the in-cell expressome.** To better resolve local density and to assess the flexibility of the 2,952 sub-tomograms, multi-body refinement implemented in RELION was employed. (**A**) The consensus refinement result was segmented into 3 bodies and correspondingly, three local masks were generated, i.e. RNAP-NusA, 30S, and 50S. (**B**) FCS curves for the 3 bodies were calculated separately. (**C**) Superposition of densities of the 3 bodies. (**D**) Top view of the RNAP density with homology model fitted. (**E**) Principal component analysis on the relative orientations of all bodies was performed to characterize the most predominant motions, represented by different eigenvectors. Relative movements corresponding to the 3 most dominant eigenvectors (red) are shown in Movie 1.

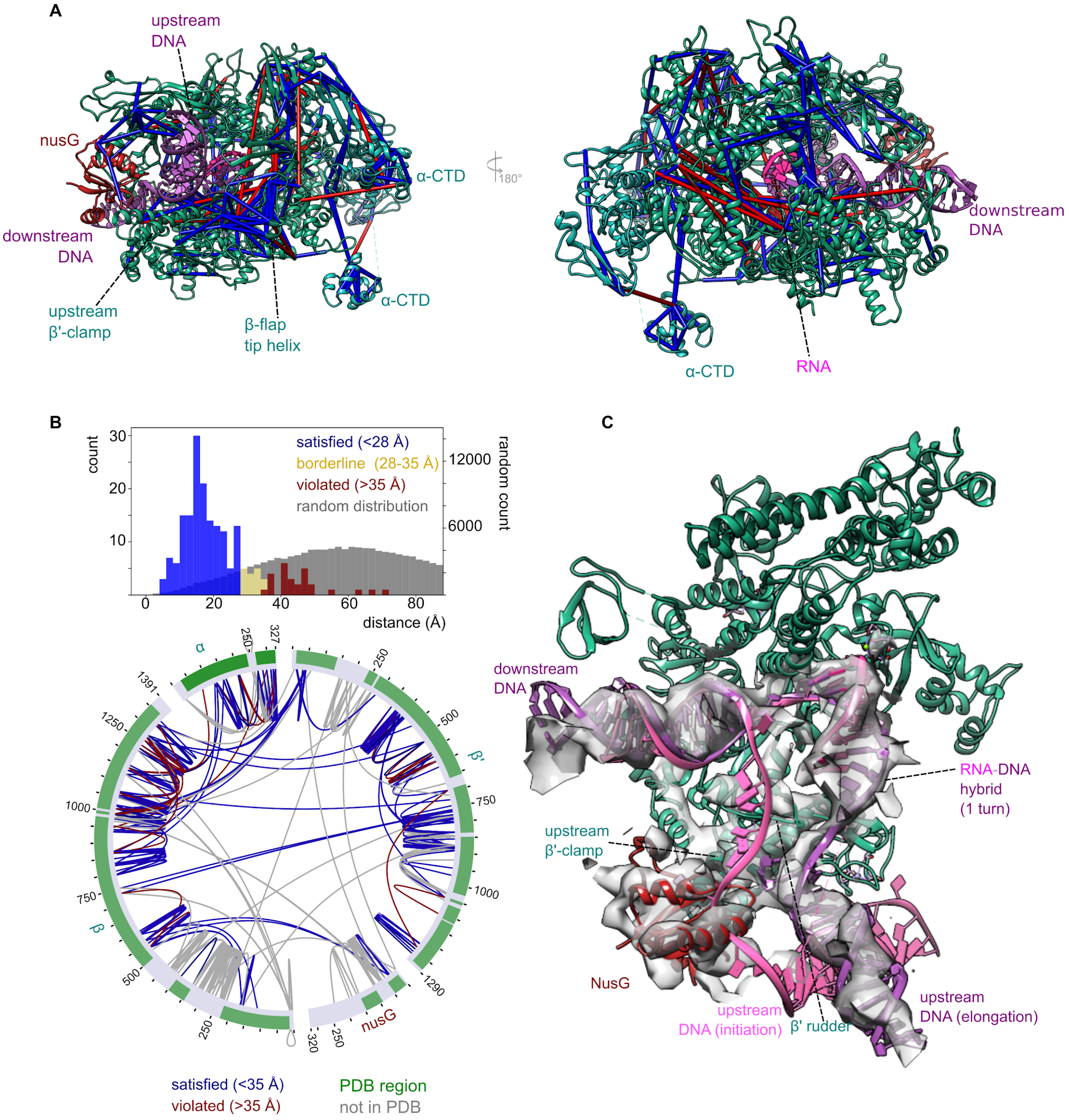


**Fig. S15.**

**The *M. pneumoniae* RNAP homology model**. (**A**) Crosslinks mapped on homology models of the core RNAP based on the *E. coli his*-paused elongation complex with nucleic acids included (PDB 6FLQ) generated using Swissmodel. The regions which were kept fully flexible in the integrative modeling or are disordered in the homology model, have been removed. The α-CTDs and β flap-tip helix are structured, but flexible with respect to the core RNAP. NusG is placed according to the EM density in the same binding mode as in the *E. coli* RNAP elongation complex (PDB 6C6U, (*69*). Satisfied links (<35 Å) are displayed in blue, and violated links (>35 Å) in red. (**B**) Top: distribution of 194 crosslink distances obtained from the RNAP model compared to a theoretical random distribution of lysine-lysine and lysine-Ntermini crosslinks from the same model. Bottom: Circle view of the crosslink distances in the model. Regions missing in the model are shaded in gray. (**C**) The β', DNA and RNA-DNA hybrid. The density in the RNAP region of the multibody-refined expressome map corresponding to the DNA, RNA-DNA hybrid and upstream β' clamp are shown. For comparison, the DNA path in the initiation RNAP holoenzyme (PDB 5VT0, (*88*) is shown in light pink. Moreover, the RNAP homology model shows a good fit to the EM density.

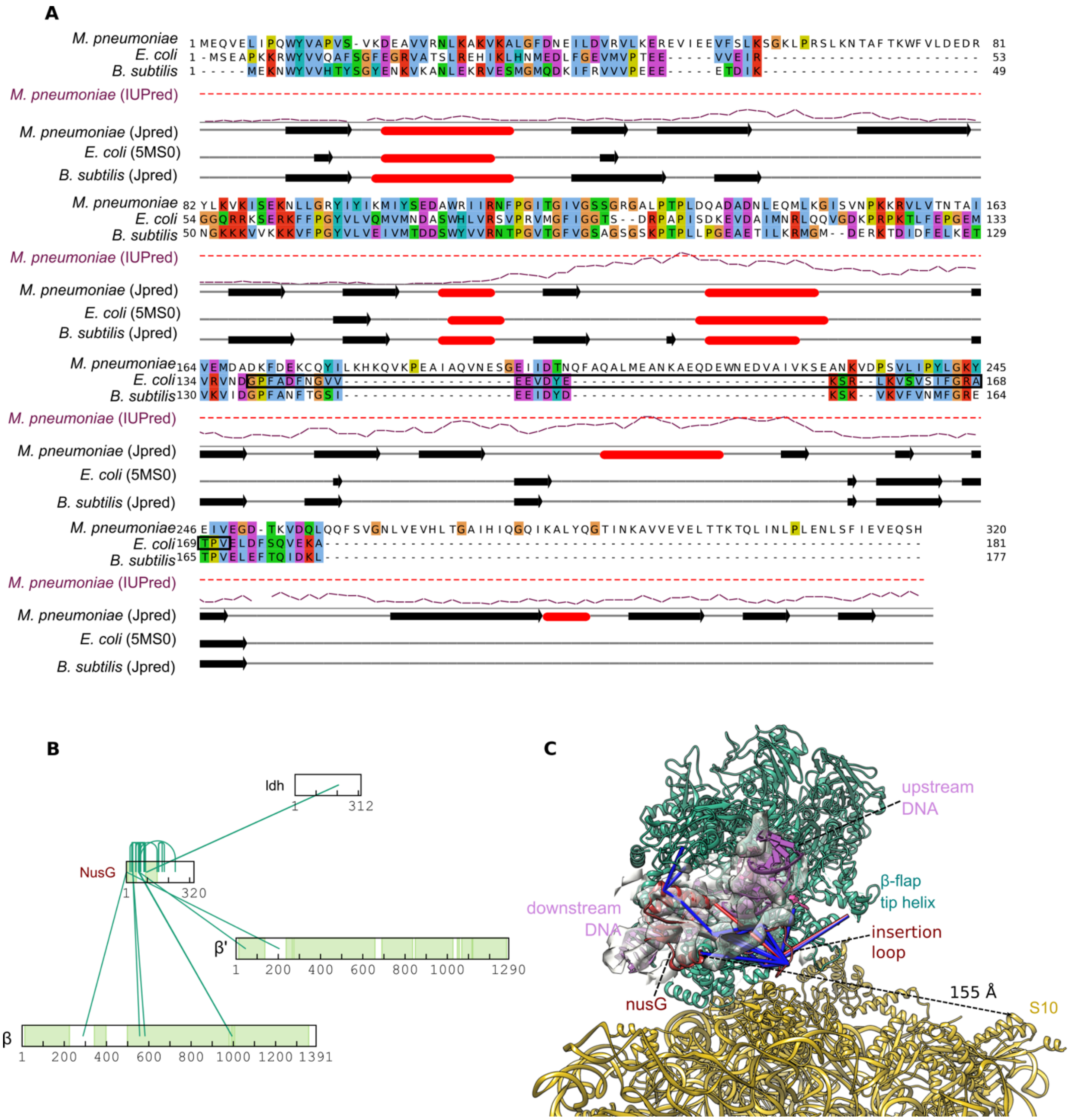

**Fig. S16.**

**NusG in *M. pneumoniae*.** (**A**) Sequence alignment of *M. pneumoniae*, *E. coli*, and *B. subtilis* NusG proteins. For *E. coli*, the secondary structure as observed in the λ phage antitermination complex (PDB 5MS0) is depicted. For *M. pneumoniae* and *B. subtilis*, the secondary structure was predicted with Jpred. β-sheets are shown as arrows, α-helices as cylinders. The long-range disorder prediction of IUPred is also shown, with the threshold of 50% disorder likelihood displayed as a dashed line. The residues involved in the *E. coli* NusG:S10 interaction, as reported in (*8*) are highlighted in the black box in the sequence. *M. pneumoniae* presents several inserts between the

NTD (residues 1-150) and the putative S10 binding region. (**B**) Crosslink interaction network of NusG. Structured regions of RNAP and NusG, not coarse grained during the modeling procedure, are shown in green. (**C**) NusG (in red) in the active expressome. Satisfied (<35 Å) crosslinks are shown in blue, violated crosslinks in red. The insertion loop in the NTD (residues 48-91) is positioned below the upstream DNA, in agreement with previous reports (*69*), and shows crosslinks to the β flat-tip helix. The distance between the end of the NTD (residue 146) and S10 on the ribosome is marked by a dashed line.

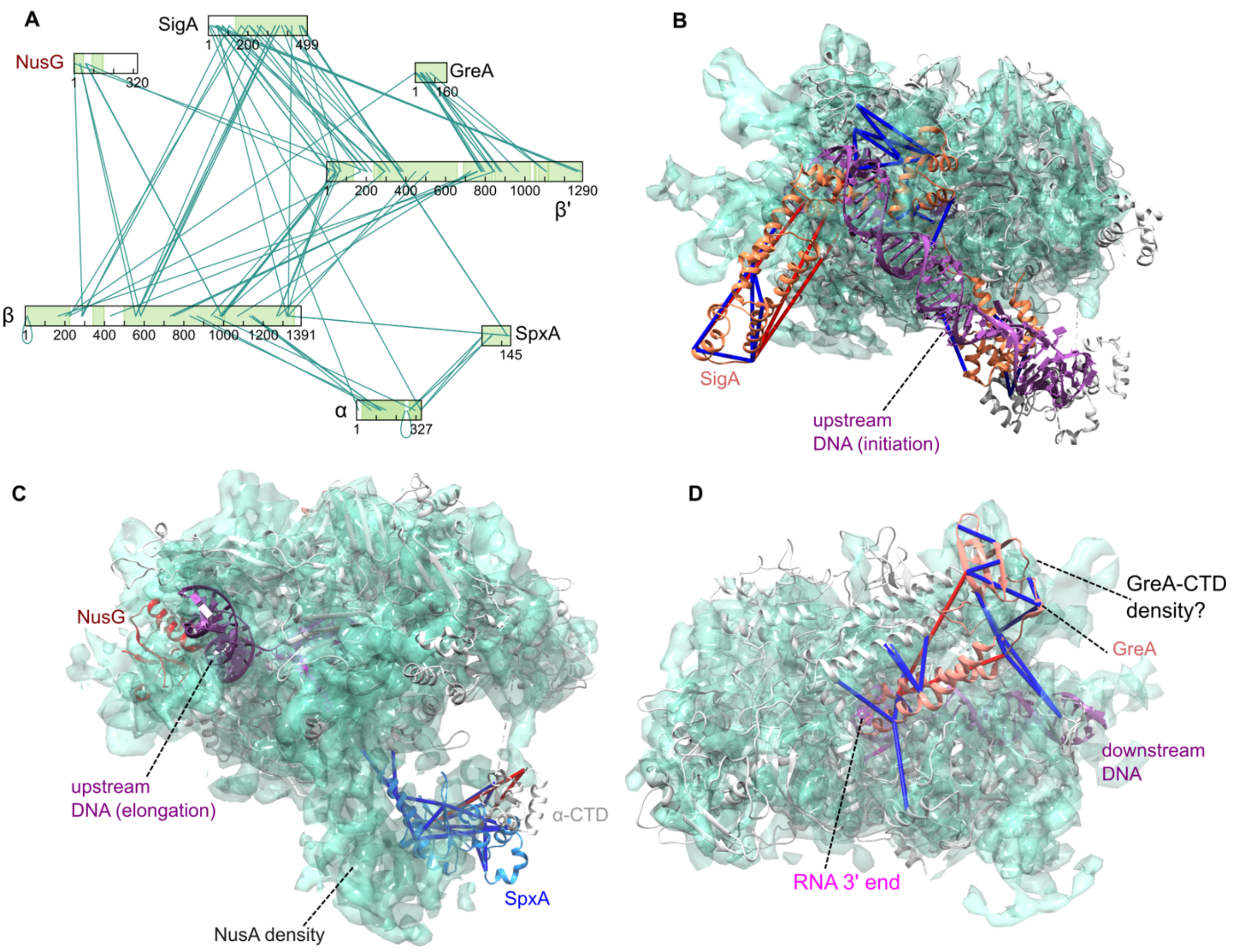


**Fig. S17.**

**Additional RNAP regulators found in the crosslink network are not constitutive components of the elongating expressome**. (**A**) Interaction network of the core RNAP α (RpoA), β (RpoB), and β' (RpoC) subunits showing the known regulators of RNAP found by in-cell crosslinking. The areas shaded in green are those present in the homology models used in this figure. (**B**) Homology model of the sigma factor SigA and DNA bound as in the RNAP holoenzyme (gray, template PDB 5VT0,(*88*)). Satisfied crosslinks (<35 Å) are shown in blue, while those longer than 35 Å are in red. The cryo-EM density assigned to RNAP is also displayed in green. The crosslinks between SigA and the core polymerase are consistent with sigma factor binding in the same mode as in the *E. coli* 70S holoenzyme, albeit with flexibility in its N-terminal region (of which residues 139-320 are present in the homology model). However, both the path of the nucleic acid and the density around the protein are not consistent with SigA being present in the expressome. (**C**) The regulatory protein SpxA, which is known to interact with the RNAP α-CTD, placed according to the *B. subtilis* model by SwissModel (template PDB 3GFK, (*89*)). It displays crosslinks to the α-CTD and the residues 1316-1362 of the β CTD. Its size, shape and position are not consistent with the expressome density. (**D**) Elongation factor GreA, responsible for resolving elongation arrest by stimulating the RNAse activity of RNAP. The binding is modelled on the structure of GreB-bound RNAP in the pre-cleavage state (PDB 6RIN, (*90*)). The crosslinks confirm the binding mode of this protein in *M. pneumoniae*. While some unassigned density could represent the GreA CTD, the lack of density around the NTD may indicate low occupancy, or a highly dynamic nature of the protein in the elongating expressome. Nevertheless, this density is filled by β-lobe 2 residues in the modeling results (Fig. S22A).

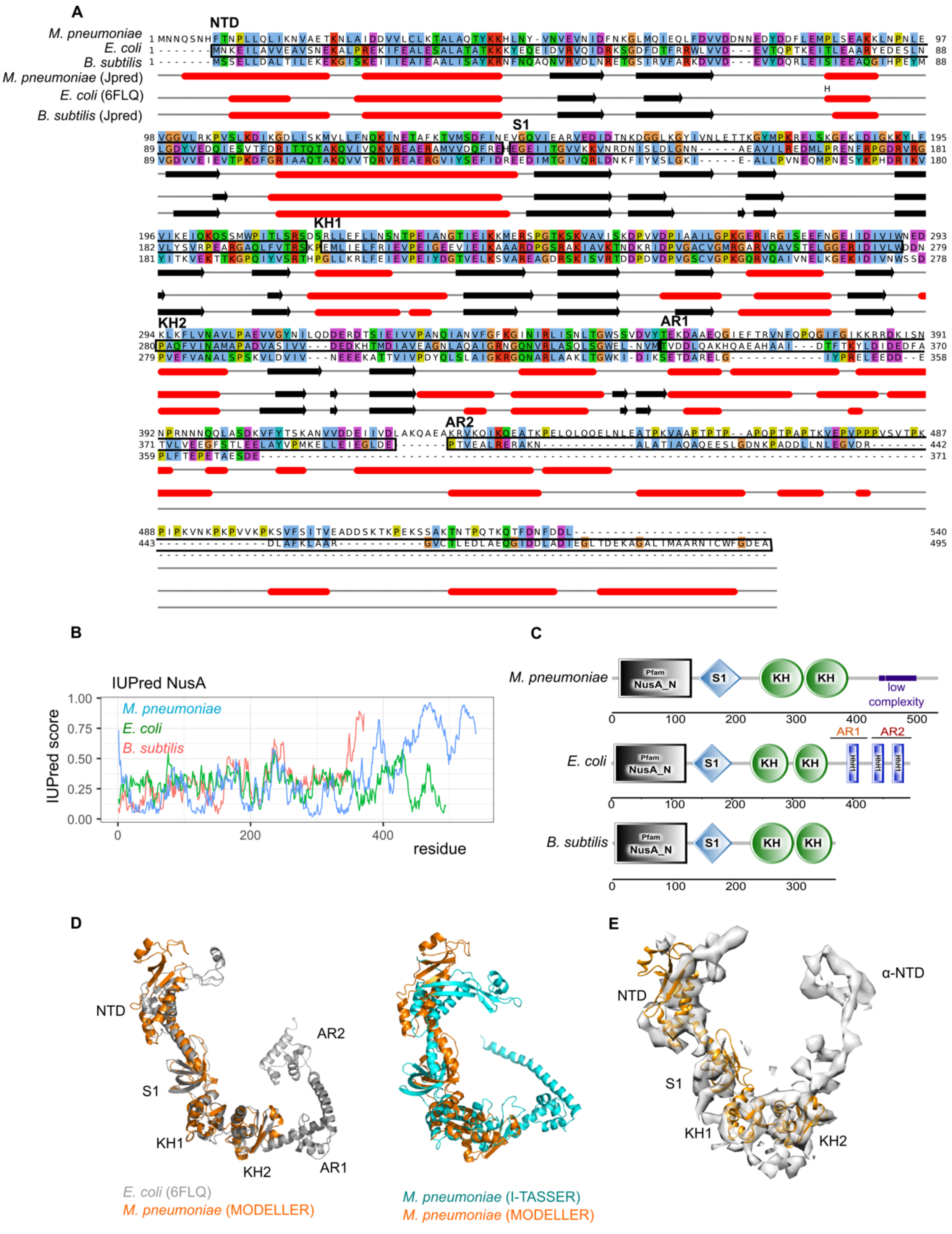


**Fig. S18.**

**NusA in *M. pneumoniae*. (A)** Sequence alignment of *M. pneumoniae*, *E. coli*, and *B. subtilis* NusA proteins. For *E. coli*, the secondary structure as observed in the RNAP paused elongation complex was used (PDB 6FLQ). For *M. pneumoniae* and *B. subtilis*, the secondary structure was predicted with Jpred. β-sheets are shown as arrows, α-helices as cylinders. The *E. coli* NusA domain boundaries are shown in black bloxes in the sequence. *M. pneumoniae* presents a highly divergent CTD, with no predicted AR2, and low sequence conservation in AR1. (**B**) Prediction of disordered regions for NusA. (**C**) The SMART database domain annotation for NusA. (**D**) Left: the *E. coli* NusA structure (gray; PDB 6FLQ) overlayed with the homology model of residues 1-364 *M. pneumoniae* NusA generated with MODELLER, used in this study (orange; rigid body boundaries in table S4). Right: homology model of NusA compared with structure prediction using I-TASSER. I-TASSER generates a structure for residues 361-457 that resembles the *E. coli* AR1 domain followed by a long extension. No fold is predicted for residues 457-540 (not displayed). (**E**) Rigid body fitting of the homology model of NusA NTD and SKK domains to the region of cryo-EM density unaccounted by RNAP. The size and shape of the density is consistent with NusA. The predicted fold generated by I-TASSER could not be fitted into the density above the second KH domain. Therefore, the NusA C-terminal region was not fitted to the EM density in integrative modeling (table S4).

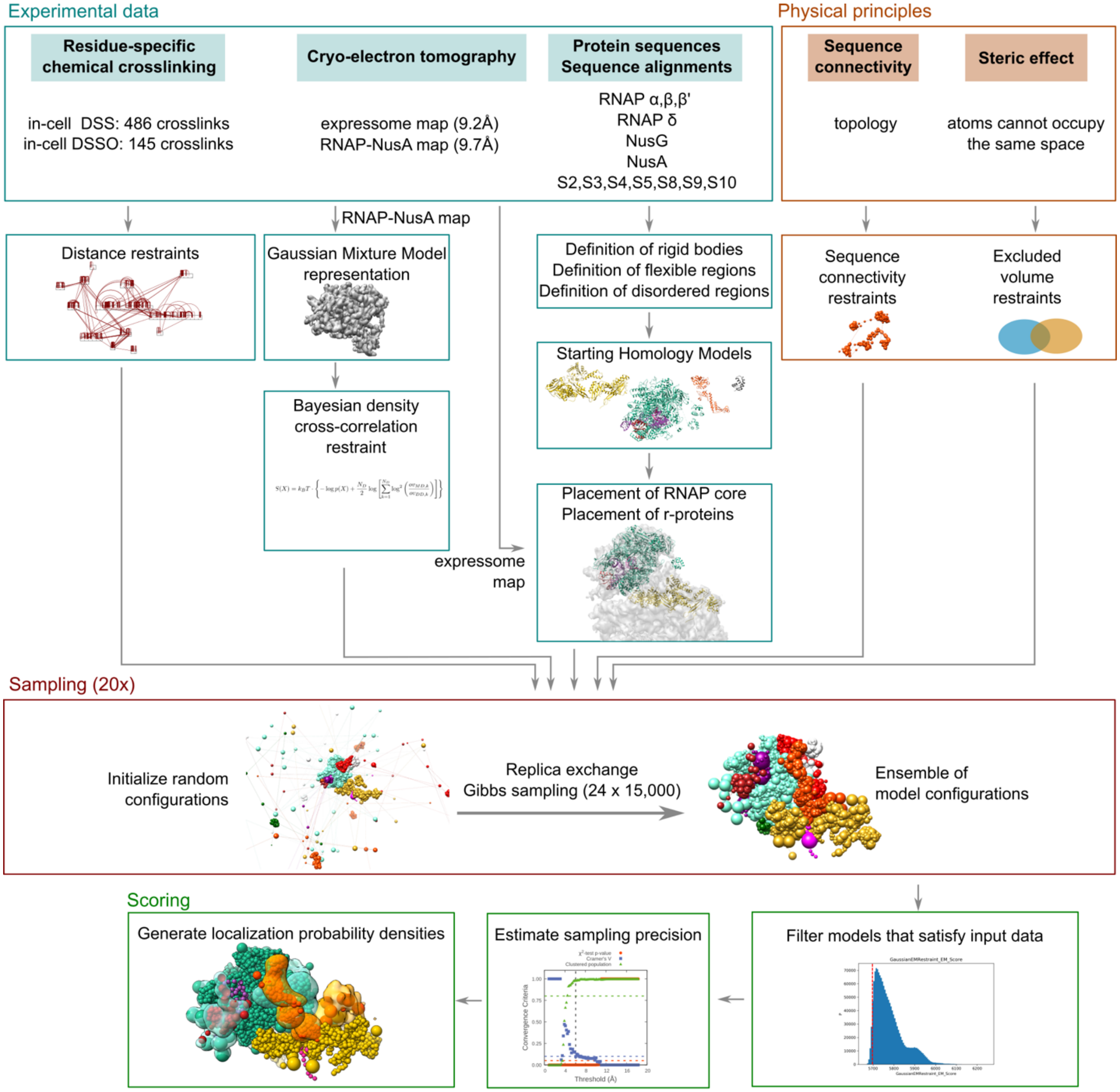


**Fig. S19.**
**Integrative modeling workflow.**

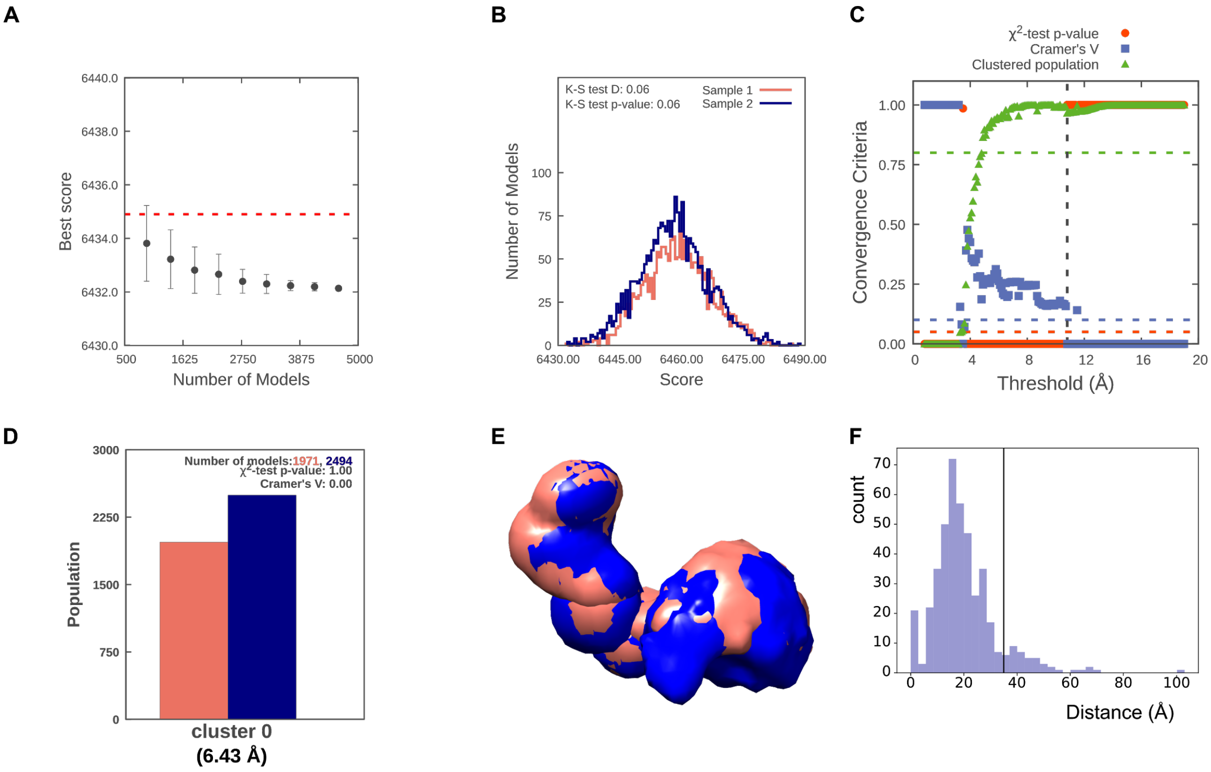


**Fig. S20.**

**Sampling completeness and precision of the integrative modeling procedure.** (**A**) Convergence of the model score in the 4623 selected models. The error bars represent the standard deviations of the scores obtained by sampling a subset of models 10 times. Score does not continue to improve as more models are sampled. (**B**) Comparison of score distributions for the selected models in even runs (Sample 1) and odd runs (Sample 2). The distributions do not differ significantly (p>0.05 in the Kolmogorov-Smirnov two-sample test) and the difference is small (D=0.06), indicating sampling completeness. (**C**) Estimation of sampling precision in the selected models as a function of rmsd sampling threshold. The vertical dashed line represents the threshold at which the p-value for homogeneity of proportions is satisfied (>0.05, the clusters significantly differ), the effect size is small (Cramer's V <0.1) and >80% of models fall within one of the selected clusters. This indicates an overall sampling precision of 10.8 Å. (**D**) Cluster populations obtained with an rmsd cutoff of 10.8 Å based on the overall sampling precision. 96% of structures fall within one cluster, with an in-cluster precision (average rmsd-based distance to the cluster centroid model) of 6.43 Å. Colors refer to models originating from sample 1 or 2 in (B). (**E**) Localisation probability densities for the NusA domains in models from the two random samples that fall within cluster 0, indicating the in-cluster variance. (**F**) Crosslink distance distribution in the cluster centroid model. The vertical line marks 35 Å. The crosslink satisfaction is 92%.

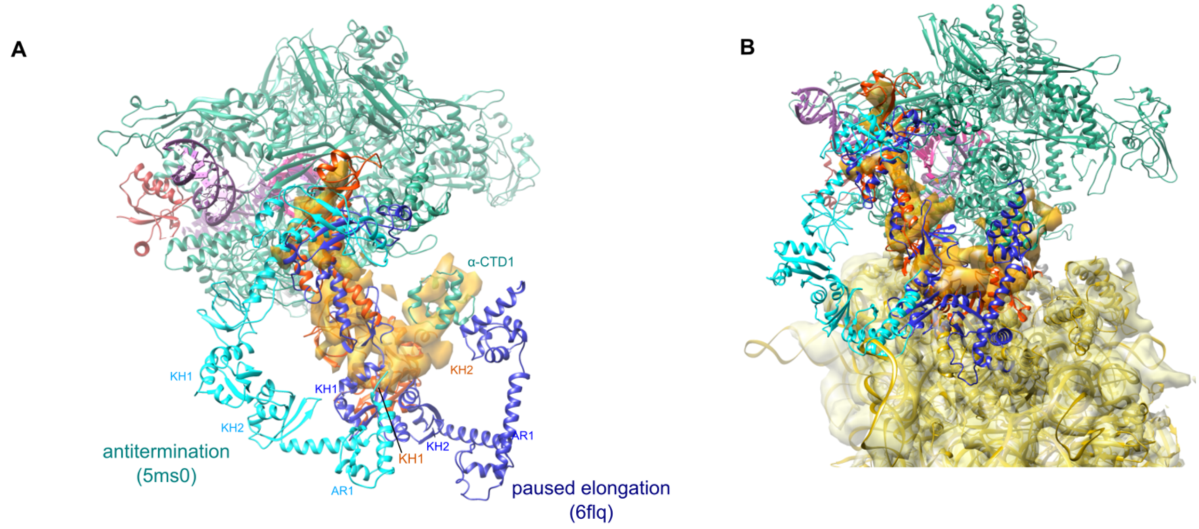


**Fig. S21.**
**NusA conformation relative to antitermination and paused elongation complexes.** (**A**) Overlay of the arrangement of NusA relative to RNAP (green) in the *M. pneumoniae* expressome, the *E. coli* λ-induced antitermination complex (cyan) and the *E. coli* paused elongation complex (dark blue). The NusA density in the multibody-refined *M. pneumoniae* elongating expressome cryo-EM ednsity is also shown (orange). While the arrangement of the NTD and S1 domains are similar to the paused elongation configuration, the KH domains are rearranged closer to RNAP in the active expressome. (**B**) The NusA arrangements as in (A) shown relative to the ribosome in the expressome structure. Both antitermination and paused elongation *E. coli* structures clash with the top of the 30S subunit. The segmented densities for the 30S subunit (yellow) and NusA (orange) are also shown.

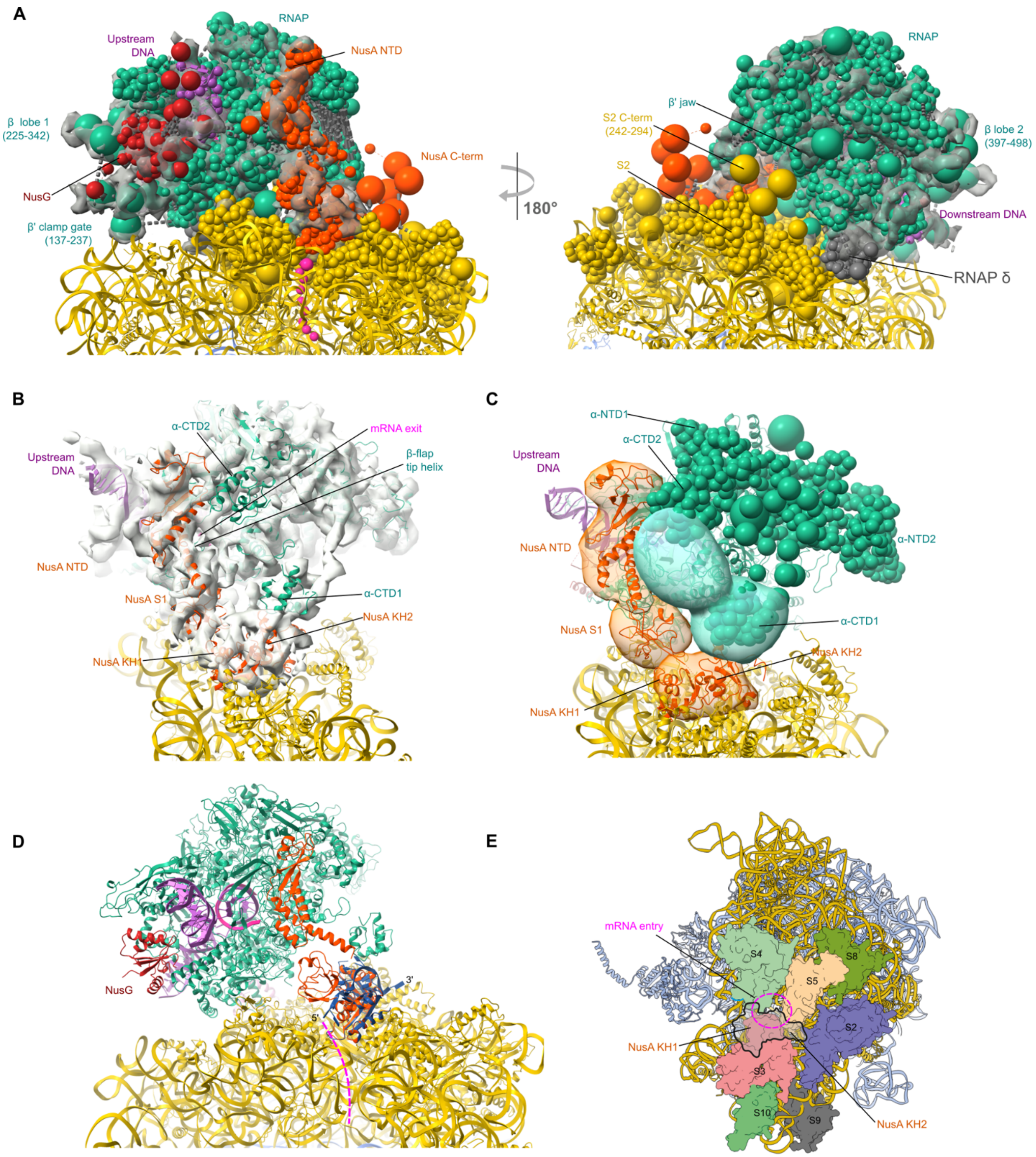


**Fig. S22.**

**Crosslinking-assisted modeling of NusA in the cryo-EM density.** (**A**) View of the centroid model of the *M. pneumoniae* elongating expressome in the multibody-refined cryo-EM map. (**B**) View of the mRNA exit channel face of RNAP with the density for NusA and the α-CTDs shown. (**C**) Localization probability densities for the NusA domains (orange) and the α-CTDs (green) displayed. The α-CTDs are localized at low precision, spanning the region between the NusA NTD and the second KH domain. (**D**) Overlay of *M. pneumoniae* NusA in the elongating expressome model (orange) with the structure of the *M. tuberculosis* NusA KH domains bound to RNA (blue) (PDB 2ATW, (*25*)). The putative mRNA path in the elongating expressome model is shown in magenta. The orientation of the mRNA on the KH domains is consistent with the RNA interacting with the KH domains near the mRNA entry site on the ribosome. (**E**) Contacts between NusA and the 30S subunit in the centroid model.

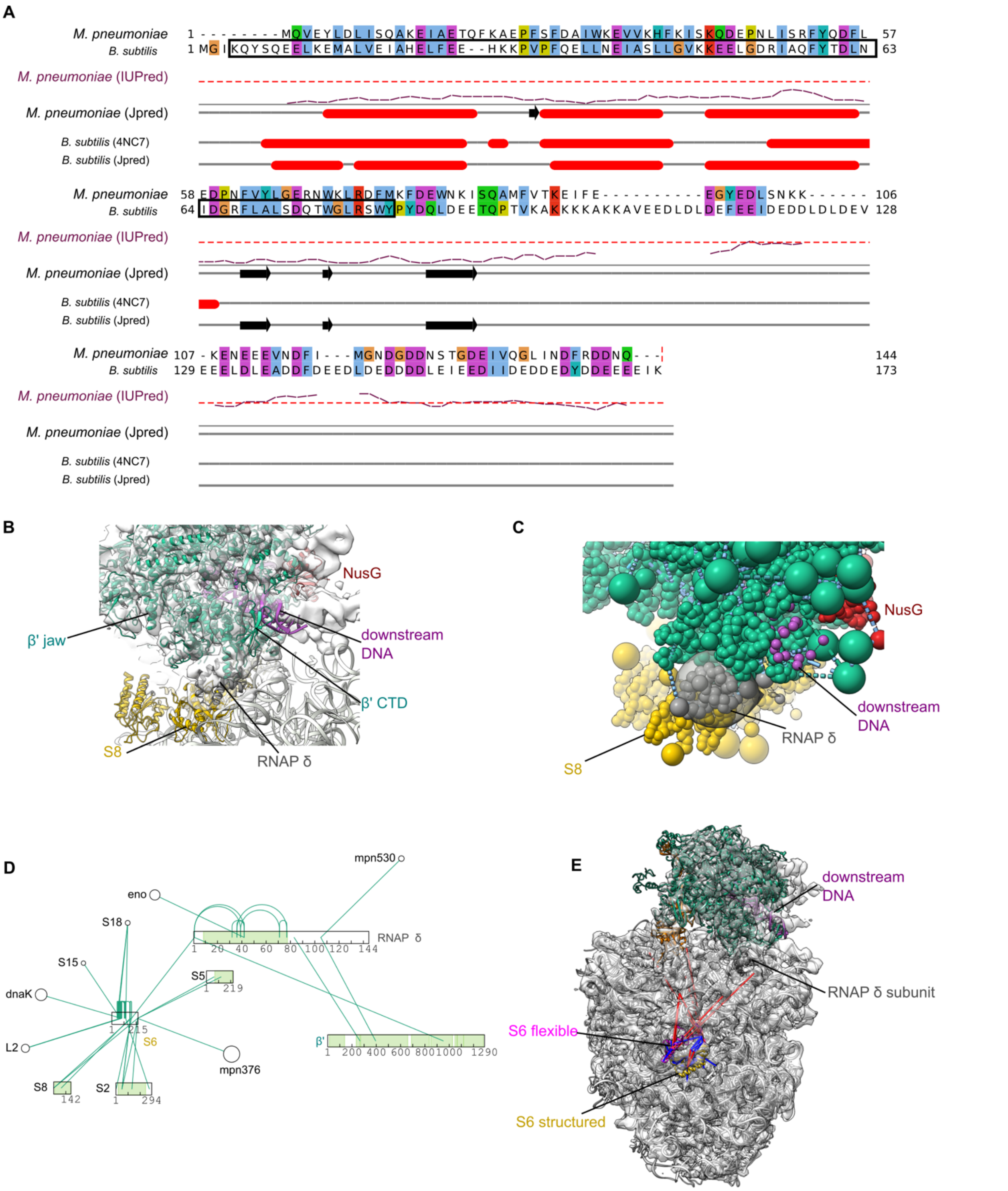


**Fig. S23.**

**Localisation of the RNAP δ subunit.** (**A**) Sequence alignment of *M. pneumoniae* and *B. subtilis* RNAP δ subunit (RpoE). The N-terminal domain, covered in the *B. subtilis* crystal structure (PDB 4NC, (*91*)) is highlighted by a black box. The secondary structure was predicted with Jpred. β-sheets are shown as arrows, α-helices as cylinders. The long-range disorder prediction of IUPred is also shown, with the threshold of 50% disorder likelihood displayed as a dashed line, indicating a disordered C-terminal region. (**B**) Placement of the NTD of the RNAP δ subunit in the elongating expressome integrative model in proximity of the downstream DNA, ribosomal protein S8 and the β' CTD. (**C**) Localisation density for RNAP δ in the most populated selected cluster displayed with crosslinks shown as dashed lines. (**D**) The crosslink interaction network of RNAP δ and ribosomal protein S6, highlighting the single link connecting the flexible RNAP δ N-terminus with the highly promiscuous S6 C-terminus. Structured regions included in the IMP protocols are shown in green. (**E**) While the structured NTD of S6 (yellow) is homologous to *B. subtilis* and is observed in the density, its C-terminal domain is likely very flexible (here, a prediction by I-TASSER is shown in magenta) and crosslinks to several 30S proteins far apart from each other (>60A). The link to RNAP δ (gray) is equally overlength (110 Å), which may be due to the disordered nature of the S6 CTD, or to the fact that this interaction occurs outside the ribosome.

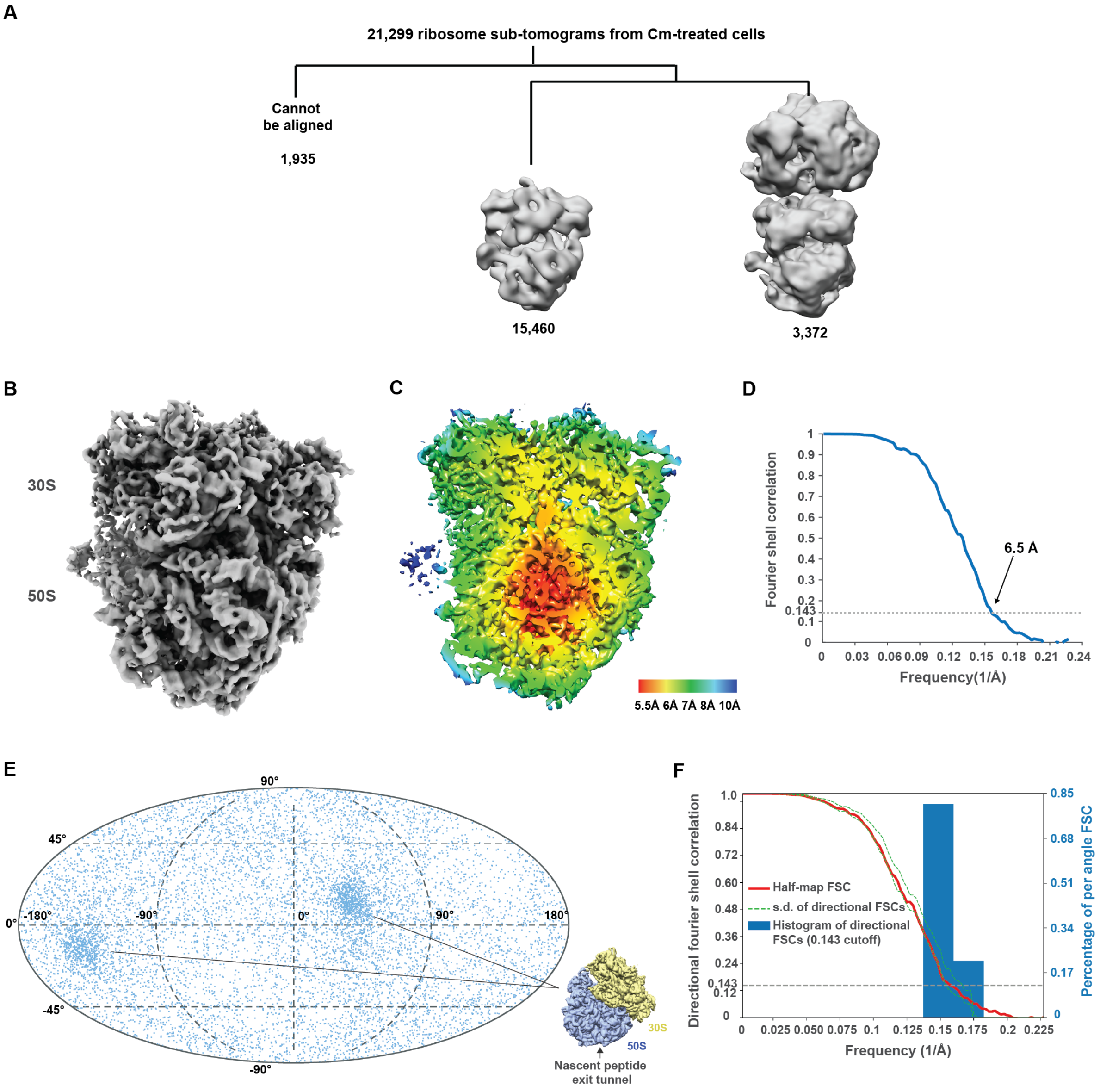


**Fig. S24.**

**Inhibiting translation by chloramphenicol de-couples RNAP and the ribosome.** (**A**) Classification of 21,299 ribosome sub-tomograms from 65 tomograms of chloramphenicol (Cm)-treated *M. pneumoniae* cells. No significant density was detected despite focused classification efforts. (**B**) Refined Cm-treated 70S ribosome density. (**C**) Central slice of the local resolution map. (**D**) Global resolution of the map is 6.5 Å (FSC=0.143). (**E**) Angular distribution map shows similar weak preferential angular distribution to the untreated dataset (Fig. S8). (**F**) 3D FSC analysis. Sphericity of 0.938 indicates low anisotropy.

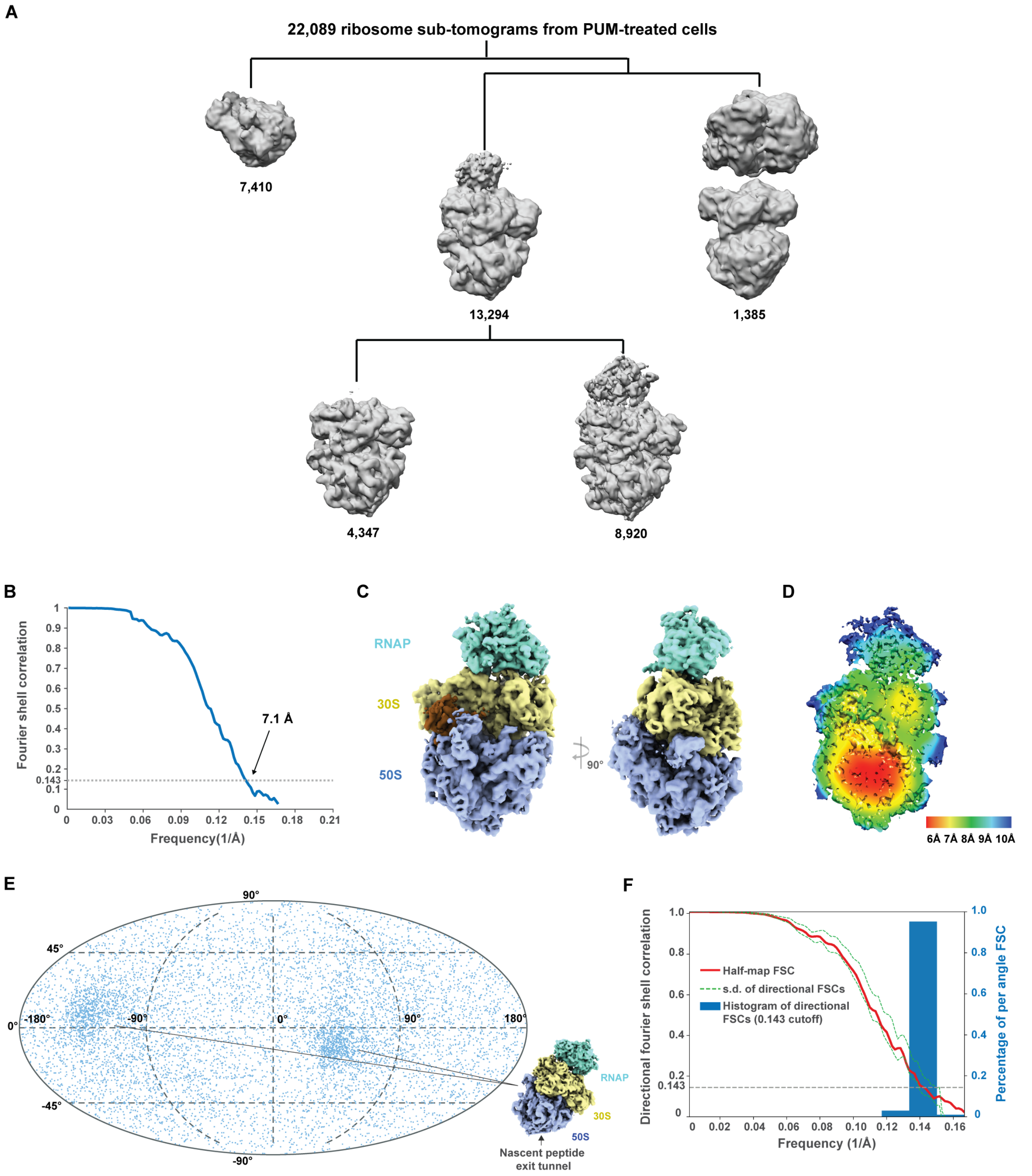


**Fig. S25.**

**Stalling RNAP by pseudouridimycin leads to rearrangement of the expressome architecture.** (**A**) Classification of 22,089 ribosome sub-tomograms from 83 tomograms of pseudouridimycin (PUM)-treated *M. pneumoniae* cells. The percentage of classified RNAP-ribosome supercomplexes increased dramatically compared to untreated cells (Fig. S7), accounting for 40% of localized ribosomes. (**B**) The supercomplex density was refined to 7.1 Å global resolution

(FSC=0.143). (**C**) Refined density of the RNAP-ribosome supercomplex. Density corresponding to NusA was not found. Identification of EF-G density (brown) associated with the ribosome indicates a pre-translocational state. (**D**) Central section of the local resolution map. (**E**) Angular distribution map of the 8,920 supercomplexes. Similar weak preferential angular distribution was observed as before (Fig. S8, 24). (**F**) 3D FSC analysis. Sphericity value of 0.942 indicates low anisotropy.

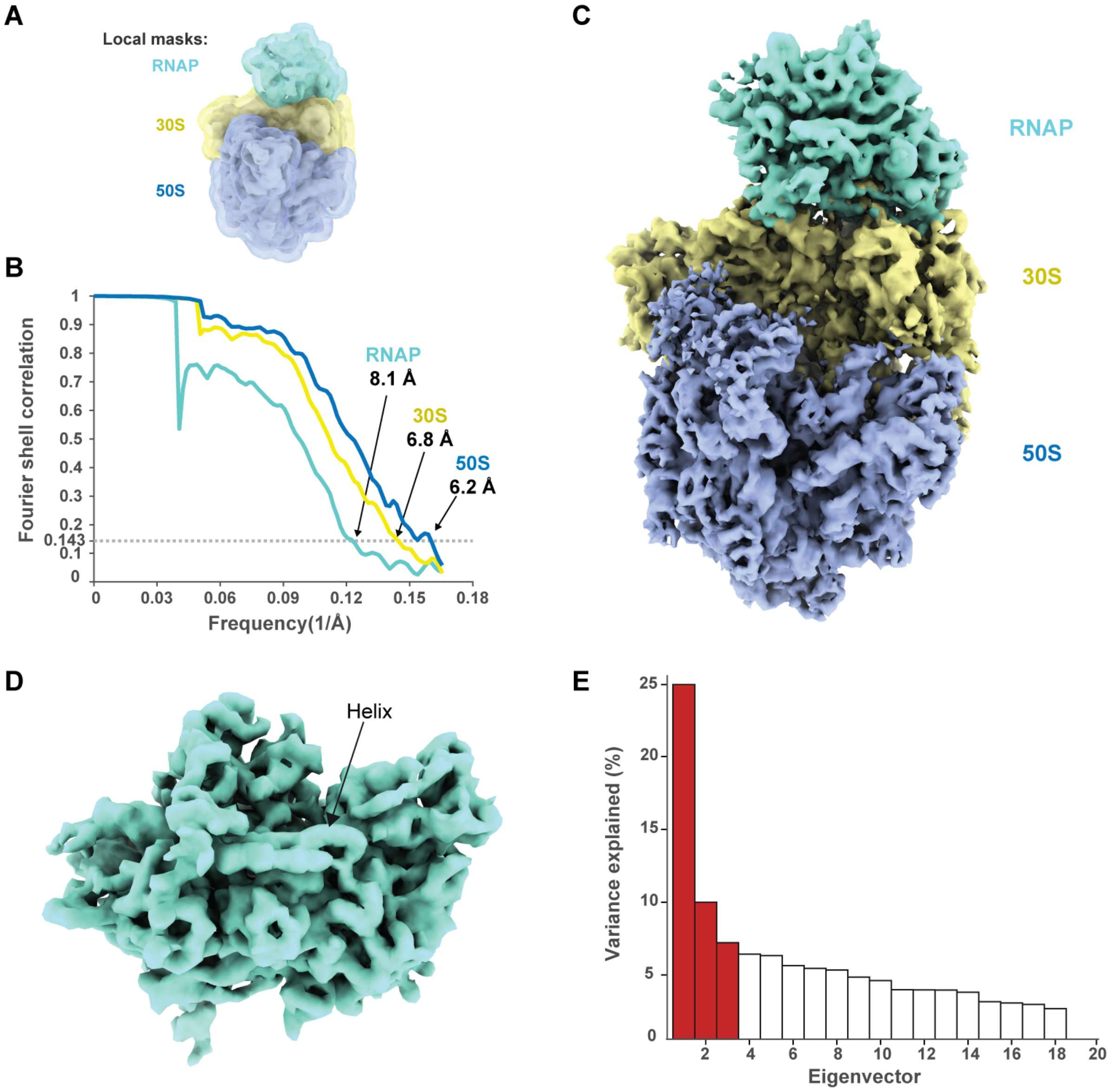


**Fig. S26.**

**Multi-body refinement of PUM-induced RNAP-ribosome supercomplexes.** (**A**) Segmented bodies from the consensus refinement and their corresponding local masks, i.e. RNAP, 30S, and 50S. (**B**) Resolution estimation for each body. (**C**) Superposition of the three bodies/densities from multi-body refinement. The global resolution of the RNAP density was estimated to be 8.1 Å (FSC=0.143), at which long helices could be clearly resolved (**D**). (**E**) Plot of variance contributions of all eigenvectors. Movements corresponding to the top three eigenvectors (in red) are shown in Movie 2.

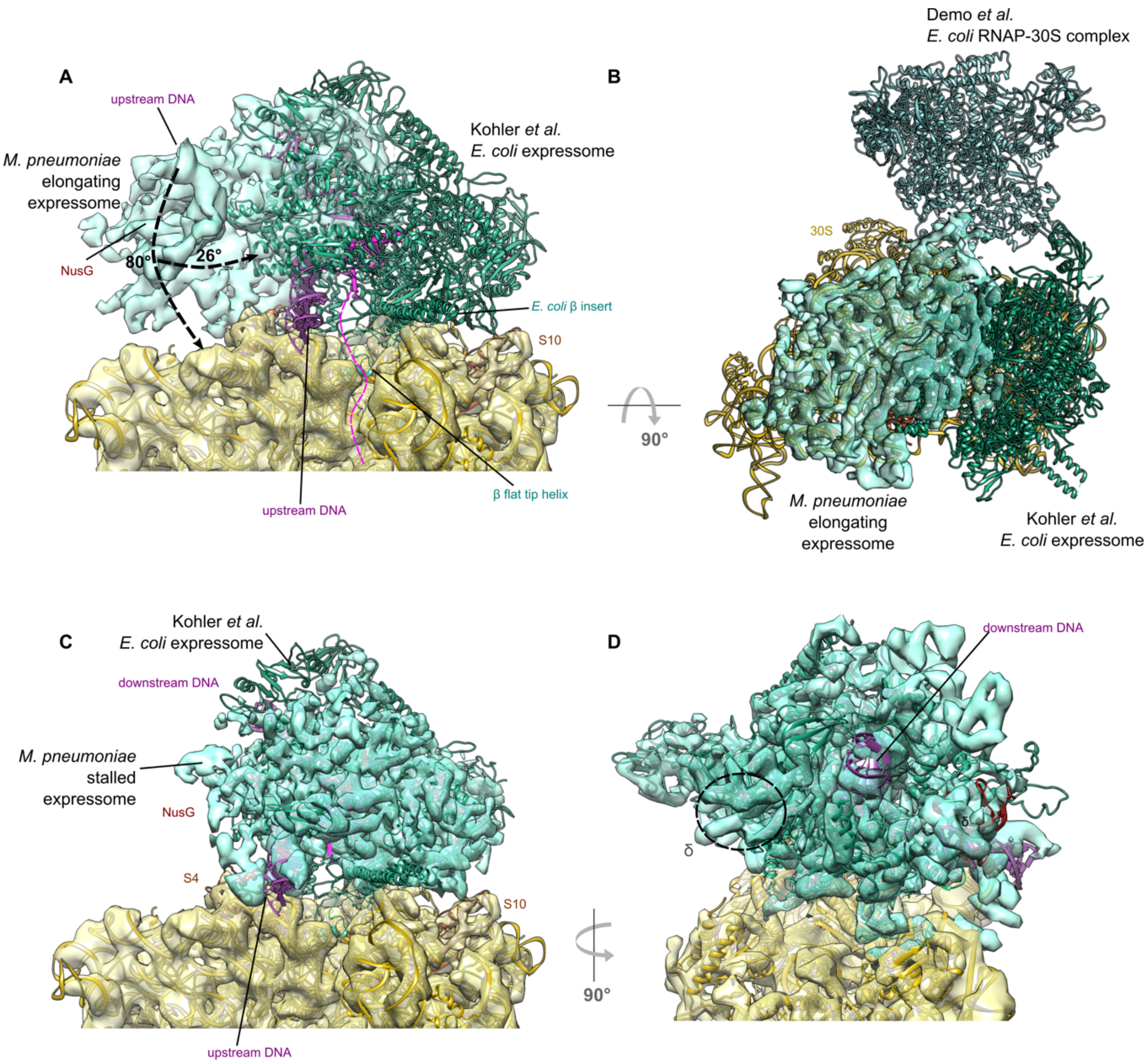


**Fig. S27.**

**Comparison of the untreated in-cell elongating expressome density and PUM-induced density to the expressome structure from Kohler *et al.*** (**A**) The untreated *M. pneumoniae* expressome densities corresponding to RNAP and the 30S subunit overlayed with the cryo-EM structure of the *E. coli* expressome in cartoon representation (PDB 5MY1, (*6*). The density corresponding to *M. pneumoniae* NusA has been omitted for clarity, and nucleic acids have been modelled in. In the *E. coli* expressome structure, the mRNA exit site on RNAP sits directly over the mRNA entry channel in the 30S, where the β flat-tip helix also resides, not leaving space for additional factors. Compared to the untreated *M. pneumoniae* RNAP orientation, RNAP tilts back towards ribosomal protein S10 and undergoes a rotation by which the upstream DNA points towards the ribosome rather than away from it. (**B**) Top view of the 30S subunit with comparison of the RNAP orientations in the untreated expressome in the present study with the orientations found in *in vitro* high resolution structures of the *E. coli* expressome (PDB 5MY1, (*6*); PDB 6AWD, (*7*). The density corresponding to *M. pneumoniae* NusA has been omitted for clarity. (**C**) The PUM-treated *M. pneumoniae* expressome densities corresponding to RNAP and the 30S subunit overlayed with the cryo-EM structure of the *E. coli* expressome in cartoon representation. Stalling *M. pneumoniae* RNAP with PUM leads to the same expressome configuration as observed in the *E. coli* expressome, indicating that this configuration may be the product of a ribosome reaching and "colliding" with a stalled RNAP. Nevertheless, *M. pneumoniae* RNAP retains NusG in the arrested state, as observed by the additional density present around the upstream DNA (Fig. 4). Interestingly, in the stalled expressome we did not detect any density corresponding to the α-CTDs. (**D**) Homology model of *M. pneumoniae* RNAP fitted into the multi-body refined RNAP cryo-EM density from PUM-treated cells. The circle highlights unexplained density at the position of δ in the elongating expressome near the β' CTD and the downstream DNA.

**Table S4. Homology models used in integrative modeling.**

| SUBUNIT (UniprotKb) | Template structure | Modeling method (quality estimates)[1] | IMP rigid bodies | Fully flexible coarse grained regions (residues per bead) |
|---|---|---|---|---|
| RNAP α/ rpoA (Q50295) | 6FLQ | SwissModel (GMQE: 0.66; QMEAN: -4.52 Seq. identity 41%)[2] | 1-233; (core)<br>234-325 (CTD) | 240-265 (5) |
| RNAP β/ rpoB (P78013) | 6FLQ | SwissModel (GMQE: 0.66; QMEAN: -4.52 Seq. identity 41%)[2] | 1-984 + 1001-1356; (core)<br>985-1000 (flat-tip helix) | 225-342; (25)<br>397-498; (25)<br>981-989; (5)<br>999-1005; (5) |
| RNAP β' /rpoC (P75271) | 6FLQ | SwissModel (GMQE: 0.66; QMEAN: -4.52 Seq. identity 41.0%)[2] | Whole sequence | 137-237; (25)<br>266-272; (25)<br>658-690; (25)<br>836-848; (25)<br>1027-1047; (25)<br>1057-1063; (25)<br>1117-1120; (25) |
| NusG 1-160 (P75049) | 6C6U | SwissModel (GMQE: 0.15<br>QMEAN: -5.91<br>Seq. identity: 22.1%) | 1-150 (NTD) | 48-91; (5)<br>147-150 (5) |
| RNAP δ/ rpoE 1-85 (P75090) | 2KRC; 2M4K; 4NC7 | MODELLER (GA341: 0.8911 z-DOPE: -1.183) | 1-85 (structured region linked to RNAP) | 81-85 (5) |
| S5/rpsE (Q50301) | 3J9W | SwissModel (GMQE 0.61 QMEAN -1.44 Seq. identity 51.6%) | Whole sequence | 1-69; (25)<br>213-219 (25) |
| S2/rpsB (P75560) | 3J9W | SwissModel (GMQE 0.60 QMEAN -5.32 Seq. identity 35.3%) | Whole sequence | 1-19; (25)<br>242-294 (25) |
| S10/rpsJ (P75581) | 3J9W | SwissModel (GMQE: 0.70 QMEAN -4.93 Seq. identity 40.6%) | Whole sequence | 1-11 (25) |
| S3/rpsC (P41205) | 3J9W | SwissModel (GMQE: 0.61 QMEAN -3.14 Seq. identity 43.1%) | Whole sequence | 207-273 (25) |
| S4/rpsD (P46775) | 3J9W | SwissModel (GMQE: 0.76 QMEAN -2.71 Seq. identity 45.7%) | Whole sequence | 203-205 (25) |
| S8/rpsH (Q50304) | 3J9W | SwissModel (GMQE 0.76 QMEAN: -1.24 Seq. identity: 50.8%) | Whole sequence | 1-9; (25) |

| | | | | |
|---|---|---|---|---|
| S9/rpsI (P75179) | 3J9W | SwissModel (GMQE: 0.78 QMEAN: -3.32 Seq.identity: 56.9%) | Whole sequence | 1-5; |
| NusA (P75591) | 1HH2;<br>1L2F;<br>5LM7;<br>6FLQ;<br>6GOV;<br>5MS0;<br>6J9E;<br>5LM9;<br>4MTN;<br>1K0R;<br>2ASB;<br>2ATW | MODELLER: (GA341 1.00 zDOPE: 0.42) | 1-140 (NTD);<br>145-213 (S1 domain);<br>214-364 (KH1 and KH2) | 1-11;<br>141-145;<br>212-216;<br>365-540 |

1: Homology model scores refer to the overall model quality prior to removal and coarse graining of any unstructured regions or regions not covered by alignment in subsequent integrative modeling steps.
2: Modelled as a single complex. Quality estimates and sequence identity refer to the combined α-β-β' model.

**Table S5.**
**Coarse graining and gaussian mixture model (GMM) representations in integrative modeling.** Each region was-coarse grained at the resolutions described. The "rigid body" column indicates whether these proteins belong to the same or different building blocks (numbering is arbitrary). Building block 5 (RNAP) and 10 (30S) were fitted as single bodies with Chimera and were not allowed to move during modeling, except for the "fully flexible regions" annotated in Table S4.

| **Protein** | **UniProtID** | **Residues** | **Residues per Bead** | **Residues per Gaussian** | **Belonging to rigid body** |
|---|---|---|---|---|---|
| NusA | P75591 | 1,142 | 5 | 10 | 1 |
| NusA | P75591 | 143,213 | 5 | 10 | 2 |
| NusA | P75591 | 214,364 | 5 | 10 | 3 |
| NusA | P75591 | 365,END | 25 | 0 | 4 |
| RNAP α | Q50295.1 | 1,233 | 25 | 10 | 5 |
| RNAP α | Q50295.1 | 234,END | 5 | 10 | 6 |
| RNAP α | Q50295.2 | 1,233 | 25 | 10 | 5 |
| RNAP α | Q50295.2 | 234,END | 5 | 0 | 7 |
| RNAP β | P78013 | 1,984 | 25 | 20 | 5 |
| RNAP β | P78013 | 985, 1000 | 5 | 5 | 8 |
| RNAP β | P78013 | 1001,END | 25 | 10 | 5 |
| RNAP β' | P75271 | 11,063 | 25 | 20 | 5 |
| RNAP β' | P75271 | 10,641,117 | 5 | 10 | 9 |
| RNAP β' | P75271 | 1118,END | 25 | 20 | 5 |
| NusG | P75049 | 1,150 | 5 | 10 | 5 |
| S5 | Q50301 | all | 25 | 0 | 10 |
| S2 | P75560 | all | 25 | 0 | 10 |
| S10 | P75581 | all | 25 | 0 | 10 |
| S3 | P41205 | all | 25 | 0 | 10 |
| S4 | P46775 | all | 25 | 0 | 10 |
| S8 | Q50304 | all | 25 | 0 | 10 |
| S9 | P75179 | all | 25 | 0 | 10 |
| RNAP δ | P75090 | 1,85 | 5 | 10 | 11 |

**Table S6. Integrative modeling score cutoffs.**

| Energy function term | Mean score* | Standard Deviation* | Selected Cutoff |
|---|---|---|---|
| DSS crosslink score | 625.7 | 13.3 | Imposed minimum 90% crosslinks <35 Å |
| DSSO crosslink score | 142.3 | 4.5 | Imposed minimum 90% crosslinks < 35 Å |
| Bayesian EM score | 5778.4 | 70.3 | 5688.0 |
| Excluded Volume | 40.8 | 16.5 | 30.0 |

* Mean score and standard deviations reported after pre-filtering to only include models with >90% crosslink satisfaction.

**Captions for Movies S1 and S2**

**Movie S1**.
Movements between the 3 bodies (i.e. RNAP-NusA, 30S, 50S) corresponding to each of the three most predominant eigenvectors, shown successively, for the native elongating expressome cryo-EM density.

**Movie S2.**
Movements between the 3 bodies (i.e. RNAP, 30S, 50S) corresponding to each of the three most predominant eigenvectors, shown successively, for the PUM-stalled expressome cryo-EM density.

36. M. C. Chambers, B. Maclean, R. Burke, D. Amodei, D. L. Ruderman, S. Neumann, L. Gatto, B. Fischer, B. Pratt, J. Egertson, K. Hoff, D. Kessner, N. Tasman, N. Shulman, B. Frewen, T. A. Baker, M.-Y. Brusniak, C. Paulse, D. Creasy, L. Flashner, K. Kani, C. Moulding, S. L. Seymour, L. M. Nuwaysir, B. Lefebvre, F. Kuhlmann, J. Roark, P. Rainer, S. Detlev, T. Hemenway, A. Huhmer, J. Langridge, B. Connolly, T. Chadick, K. Holly, J. Eckels, E. W. Deutsch, R. L. Moritz, J. E. Katz, D. B. Agus, M. MacCoss, D. L. Tabb, P.

Mallick, A cross-platform toolkit for mass spectrometry and proteomics. *Nat. Biotechnol.* **30**, 918–920 (2012).

79. D. Mellacheruvu, Z. Wright, A. L. Couzens, J.-P. Lambert, N. A. St-Denis, T. Li, Y. V. Miteva, S. Hauri, M. E. Sardiu, T. Y. Low, V. A. Halim, R. D. Bagshaw, N. C. Hubner, A. Al-Hakim, A. Bouchard, D. Faubert, D. Fermin, W. H. Dunham, M. Goudreault, Z.-Y. Lin, B. G. Badillo, T. Pawson, D. Durocher, B. Coulombe, R. Aebersold, G. Superti-Furga, J. Colinge, A. J. R. Heck, H. Choi, M. Gstaiger, S. Mohammed, I. M. Cristea, K. L. Bennett, M. P. Washburn, B. Raught, R. M. Ewing, A.-C. Gingras, A. I. Nesvizhskii, The CRAPome: a contaminant repository for affinity purification-mass spectrometry data. *Nat. Methods*. **10**, 730–736 (2013).
